# scprocess: a pipeline for processing, integrating and visualising atlas-scale single cell data

**DOI:** 10.64898/2026.03.09.710141

**Authors:** Maruša Koderman, Julia Pilarski, Erica Bianco, Danilo González, Mark D Robinson, Will Macnair

**Affiliations:** Department of Molecular Life Sciences, University of Zurich, Zurich, Switzerland; Department of Biosystems Science and Engineering, ETH Zurich, Basel, Switzerland; Roche Pharma Research and Early Development (pRED), Roche Innovation Center Basel, F. Hoffmann-La Roche Ltd, Basel, Switzerland; Computational Sciences Center of Excellence, F Hoffmann-La Roche Ltd, Basel, Switzerland; SIB Swiss Institute of Bioinformatics, University of Zurich, Zurich, Switzerland

## Abstract

**Motivation:** The transition toward “atlas-scale” single cell research has resulted in datasets comprising millions of cells across hundreds of samples, creating significant challenges for data management, computational efficiency, and reproducibility. While numerous methods are available for individual steps in single cell data processing, the highly complex nature of the analysis makes it challenging to maintain a clear record of every tool and parameter used. This makes final results difficult to reproduce, highlighting the need for a unified workflow that integrates multiple steps into a cohesive framework.

**Results:** *scprocess* is a *Snakemake* pipeline designed to streamline and automate the complex steps involved in processing single cell RNA sequencing data. Specifically optimized for data generated using the 10x Genomics technology, it provides a comprehensive solution that transforms raw sequencing files into standardized outputs suitable for a variety of downstream tasks. The pipeline is built to support the analysis of datasets comprising multiple (e.g. 100+) samples via a simple CLI, allowing researchers to efficiently explore their datasets while ensuring reproducibility and scalability in their workflows.

**Availability and implementation:** *scprocess* can be installed via GitHub (https://github.com/marusakod/scprocess) under the MIT license. Documentation, including setup instructions and tutorials on example datasets is available at https://marusakod.github.io/scprocess/.

## Introduction

Single cell RNA sequencing (scRNA-seq) has revolutionized biological research by enabling the transcriptomic profiling of individual cells, thereby unveiling cellular heterogeneity, rare cell populations, and dynamic cellular processes that are obscured by traditional bulk RNA sequencing methods. Rapid maturation of these technologies and a significant reduction of per-cell sequencing costs has resulted in an exponential increase in data generation (Svensson et al. 2018). The most common are platforms that leverage droplet-based microfluidics to partition cells for cDNA synthesis and molecular barcoding. By providing the throughput necessary to capture thousands of cells per experiment, these methods have enabled the construction of so-called single cell “atlases” comprising millions of cells across diverse cohorts (Sikkema et al. 2023; Gabitto et al. 2024; Siletti et al. 2023; Yao et al. 2023; Jiménez-Gracia et al. 2026). This deluge of data presents considerable computational challenges, including data storage, processing capabilities, and the need for specialized algorithms to extract meaningful biological information. Although many methods have been developed that address specific steps in scRNA-seq data analysis, researchers must nevertheless navigate dozens of critical decision points throughout the pipeline. This high degree of methodological flexibility makes the transition from raw data to the final output highly sensitive to user choices, posing significant challenges to reproducibility and highlighting the need for transparent, reproducible workflows.

In this paper we present *scprocess* which consolidates diverse processing tools for scRNAseq-data analysis into a unified framework. In the following sections we present an overview of the pipeline’s modular structure and discuss the principles guiding our implementation choices. We provide a functional comparison of *scprocess* with other existing tools in Supplementary Table 1. Tutorials, usage guidelines, references for all parameters and other additional details can be found in the accompanying documentation.

### Pipeline overview

*scprocess* is a *Snakemake*-based pipeline for processing scRNA-seq and single nuclei RNA sequencing (snRNA-seq) data generated using the 10x Genomics droplet based sequencing technology (Figure 1). It aims to facilitate execution of common preprocessing steps in single cell data analysis, irrespective of experimental design, and generates outputs suitable for a variety of downstream analyses. A key feature of *scprocess* is its ability to efficiently handle large multi-sample datasets which are becoming increasingly common in single cell research. These datasets often require high-performance computing (HPC) systems due to their size and complexity. To meet these demands, *scprocess* is specifically optimized for HPC environments, employing various strategies for memory management and runtime optimization. *scprocess* leverages conda environments to manage dependencies, ensuring the analyses are reproducible across different systems.

**Figure 1:**
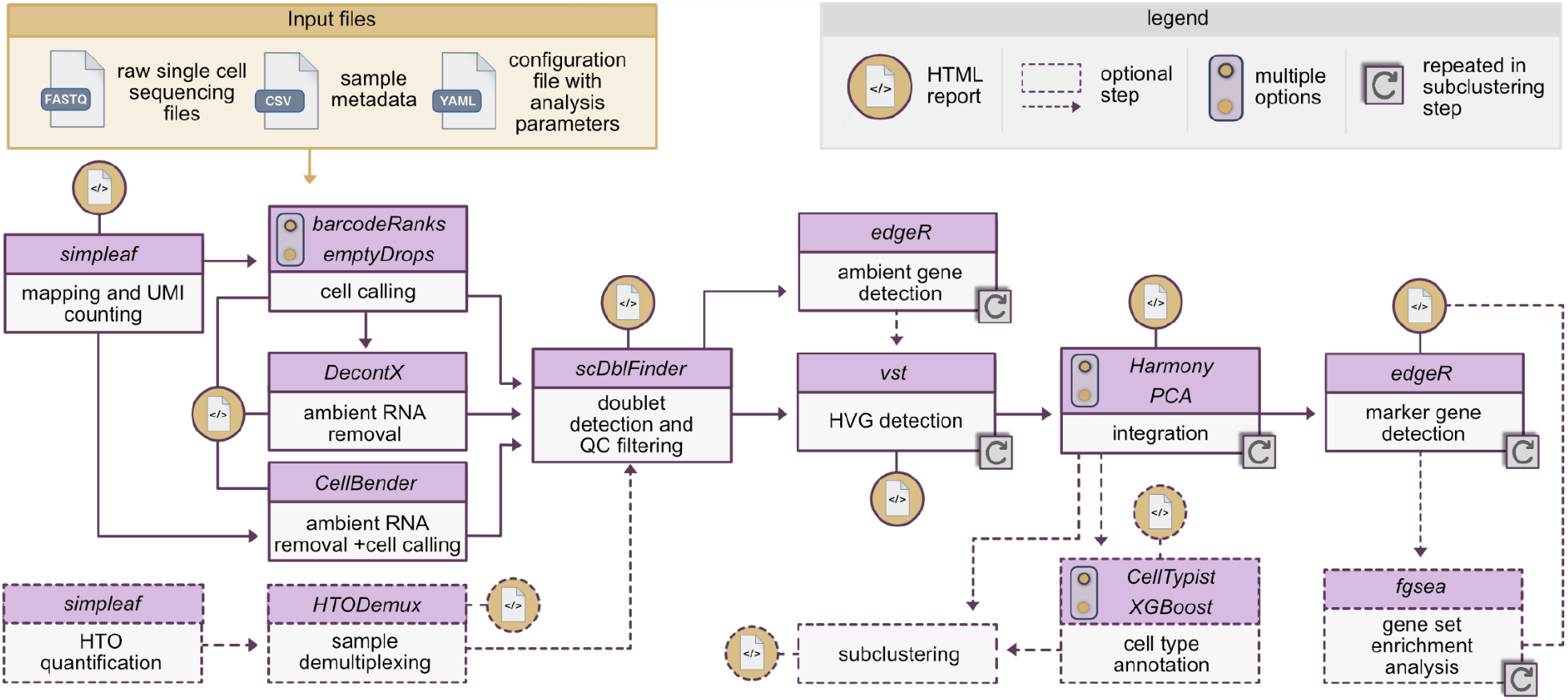
Overview of *scprocess* including input file requirements, all pipeline steps and specific software packages/algorithms used. HVG: highly variable gene; HTO: hashtag oligos.

Single cell data analysis often requires users to iterate though different parameters and make adjustments after inspecting the results. To support this, *scprocess* is modular, allowing users to execute individual steps independently. This feature, combined with detailed HTML reports generated for all major pipeline steps, enables users to assess intermediate outputs, refine parameters based on the outputs, and rerun previous steps if necessary. Reports generated using *scprocess* on a dataset comprising 149 samples (Pineda et al. 2024) are available as Supplementary File 1 and at https://marusakod.github.io/scprocess_pineda_2024/.

The configuration of *scprocess* is managed via a YAML file that defines the analysis parameters and serves as one of the three main inputs to the pipeline, alongside raw sequencing files and sample metadata. The YAML file can act as a record of what parameters were used to process a dataset.

### Read alignment and quantification

*scprocess* takes FASTQ files as input and, in the first step, performs alignment of sequencing reads to a reference genome, and quantification of unique molecular identifiers (UMIs). *Cell Ranger*, the standard tool for this task, relies on the *STAR* genome-based aligner (Dobin et al. 2013) and is notably resource-intensive (He et al. 2022). For datasets comprising hundreds of samples, slow processing and high memory use can become serious limiting factors, making alignment potentially one of the most costly steps in a workflow. To address these scalability challenges, *scprocess* uses *simpleaf*, a tool which facilitates access to features of the *alevin-fry* ecosystem (He and Patro 2023). *Alevin-fry* is a pseudoaligner that offers a significantly faster and more memory-efficient alternative to genome-based alignment tools and generates count matrices with separate spliced and unspliced read counts for each gene (He et al. 2022). This is particularly useful for various downstream tasks such as quality control and RNA velocity analysis. Notably, *scprocess* includes functionality for automatic detection of 10x Genomics assay chemistry, a feature absent in *simpleaf*.

### Cell calling and ambient RNA removal

In droplet-based platforms, only a small proportion of droplets contains an intact cell (Thompson et al. 2014). Meanwhile, cell-free or ambient mRNA released during sample preparation (often due to cell lysis or leakage) is encapsulated in nearly all droplets, regardless of their cellular content; most droplets therefore contain ambient RNA only. Distinguishing between the two droplet populations is typically automated and serves as a prerequisite for further processing. Additionally, correcting for ambient RNA can help minimize distortions in expression estimates and downstream analyses (Caglayan et al. 2022; Arora et al. 2025).

*scprocess* provides two primary options for cell calling and ambient RNA removal, allowing users to choose the method best suited to their preferences and hardware availability. The first option, *CellBender*, models the profile of empty droplets to perform both cell calling and ambient RNA removal (Fleming et al. 2023). While a recent benchmark identified this approach as providing the most precise estimates of background noise (Janssen et al. 2023), it is highly resource-intensive, often requiring over 3 hours to process a single sample, even with GPU acceleration. For users without GPU access or those aiming for greater processing efficiency, *scprocess* also offers *DecontX*, a CPU-based alternative that processes individual samples in minutes (Yang et al. 2020). Since DecontX does not perform cell calling, *scprocess* enables users to first identify cells using an approach based on the *barcodeRanks* function from the *DropletUtils* package (Lun et al. 2019; Griffiths et al. 2018), which separates droplet populations by detecting key transition points on the barcode-rank curve, or the *emptyDrops* approach, which tests barcodes for significant deviations from the ambient profile (Lun et al. 2019). *scprocess* calls *DecontX* with an estimated ambient RNA profile as input (rather than the *DecontX* default that uses a cluster-based background model); this approach was shown to provide more accurate noise estimations (Janssen et al. 2023).

### Quality control

A critical step in sc/snRNA-seq workflows is quality control (QC), which aims to exclude suboptimal capture events and cellular debris while preserving informative data and preventing biases in cell type retention. *scprocess* allows filtering based on library size (number of UMIs), number of detected features, mitochondrial read proportion and proportion of spliced reads per barcode. With the exception of spliced proportion, these metrics are the standard in the field, however identifying a distinct “poor quality” cell population based on them is often difficult. For instance, applying stringent library size and feature counts thresholds may exclude real cell populations with lower RNA content (Macnair and Robinson 2023; Hippen et al. 2021).

The interpretation of mitochondrial content requires particular caution in single nucleus experiments. Filtering is traditionally based on the assumption that properly isolated nuclei should be devoid of cytoplasmic organelles. However, research indicates that in response to stress, the nucleus can form physical contact sites with mitochondria (Desai et al. 2020), and consequently, isolating nuclei may inadvertently capture mitochondria, especially in cells from diseased samples. In such cases, aggressive filtering based on mitochondrial proportions could disproportionately exclude populations of interest.

A relatively novel QC metric is the percentage of spliced or intronic reads. Because mature mRNA is primarily located in the cytoplasm, a high proportion of spliced reads in a single nucleus library often indicates significant cytoplasmic contamination (Macnair and Robinson 2023; Montserrat-Ayuso and Esteve-Codina 2024). In addition to having a strong biological rationale, this metric consistently exhibits a bimodal distribution in snRNA-seq data, providing a clearer basis for excluding droplets that likely contain cytoplasmic material.

It is common practice to calculate sample-specific QC thresholds using the median absolute deviation (MAD), however this approach can be problematic in samples composed of multiple cell types with divergent distributions of QC metrics (Macnair and Robinson 2023). *scprocess* therefore relies on user-specified thresholds, that can be defined for individual samples and are ideally informed by inspecting the metric distributions.

In contrast to metrics such as read and feature counts, where defining “poor quality” is somewhat subjective due to a lack of empirical ground truth, doublet detection methods can be validated using datasets with experimentally annotated doublets. A comprehensive benchmark of these methods demonstrated that *scDblFinder* outperforms other tools in identifying multiplets (Germain et al. 2021; Xi and Li 2021a, 2021b) and has therefore been included into *scprocess*.

### Selection of variable features

Following QC on individual samples, analysis typically shifts to an integrated dataset containing the full set of genes and cells. However, for datasets comprising a large number of samples, concatenating unfiltered count matrices can become highly memory-intensive, potentially even to the limits of the number of non-zero entries in a matrix (2^31-1). Principal component analysis (PCA) - which is required for clustering and uniform manifold approximation and projection (UMAP) - is however generally not performed using the full set of features. Instead, a subset of features that capture meaningful variability within the dataset is selected, to improve computational efficiency and prioritize biological signal over technical noise.

*scprocess* facilitates the analysis of large datasets by eliminating the need to load the entire concatenated count matrix into memory for selection of highly variable genes (HVGs). It achieves this by applying a slight modification of the *Scanpy* implementation of the *Seurat vst* HVG detection method (Stuart et al. 2019) that operates on chunked count matrices. Briefly, the *vst* algorithm computes the mean and variance for each gene, models the relationship between the log-mean and log-variance using local polynomial regression (loess), and calculates a regularized variance based on the differences between the observed and smoothed variances. Naively, this requires a full matrix of genes and cells, which is impractical for large datasets. *scprocess* processes the data in chunks of genes, enabling the *vst* calculations to be performed in two sequential passes. While a similar approach has been recently proposed (Hu et al. 2025), *scprocess* provides unique flexibility by allowing these metrics to be computed independently per sample (default) or per sample group before generating a consolidated final gene ranking (following the principles in Seurat’s *SelectIntegrationFeatures* function). This approach is frequently adopted to prevent the results from being driven by sample-specific or batch-specific effects.

The selection of HVGs can significantly impact downstream analysis, yet how to determine the optimal number of genes and the most appropriate selection method are not yet known. In *scprocess*, diagnosis of feature selection focuses on the impact of ambient RNA contamination. *scprocess* aggregates counts from both cell-containing and empty droplets into two pseudobulk profiles per sample, then performs differential expression analysis using edgeR (Chen et al. 2025; Robinson et al. 2010) to identify genes enriched in empty droplets (i.e., ambient genes). Because high standardized variance in ambient genes may arise from residual contamination or variations in ambient RNA removal efficiency across samples, *scprocess* filters these features out by default during the ranking process.

### Integration

Following feature selection, the data is projected into a lower-dimensional space for clustering and visualization - a process that must also account for batch effects to prevent technical variation from confounding the results. In *scprocess* the integration step covers PCA, batch correction with Harmony (Korsunsky et al. 2019), nearest-neighbor graph construction, Leiden clustering and UMAP. On large-scale datasets, these steps have some of the longest execution times in the workflow. To address this, *scprocess* offers two alternative processing frameworks: a standard *Scanpy*-based workflow (Wolf et al. 2018), and a version with equivalent functionality using *RAPIDS-singlecell*. The latter leverages GPU acceleration to achieve massive performance boosts, particularly for clustering and UMAP (Dicks 2023; Nolet et al. 2022).

In *scprocess*, the integration step is executed twice. The initial round includes all QC-ed cells as well as those identified as doublets by *scDblFinder*. This preliminary integration allows for identification of doublet-enriched clusters, which can optionally be removed before the second round of integration including only filtered data.

### Marker gene identification

Assigning meaningful labels to clusters is essential for interpretation of single cell data. This is commonly achieved by examining marker genes for each cluster, identified by comparing the expression profile of each cluster against all others. For this task, many popular tools employ the Wilcoxon rank-sum test where each cell is treated as an independent biological replicate - an assumption that ignores the inherent dependency of cells originating from the same sample or organism. This approach has been shown to identify hundreds of differentially expressed genes even in the absence of true biological differences (Squair et al. 2021). Instead, *scprocess* employs a pseudobulk approach for marker gene identification, where aggregated counts are generated within each cluster at the sample level and compared using *edgeR* (Chen et al. 2025). While pseudobulk methods are typically reserved for comparison of a single cell type across different experimental conditions (Crowell et al. 2020; Squair et al. 2021), *scprocess* applies this logic to marker gene identification to increase statistical robustness.

To further facilitate cell type annotation, *scprocess* includes an option to perform gene set enrichment analysis (GSEA) (Subramanian et al. 2005) on the set of identified marker genes using the *fgsea* R package (Korotkevich et al. 2016). By identifying biological processes and pathways unique to each cluster, GSEA can provide additional evidence for characterizing cellular identity.

### Cell type annotation

In recent years, several large-scale expert-curated single cell datasets have been generated for a variety of tissues and organisms. (Yao et al. 2023; Braun et al. 2023; Sikkema et al. 2023; Lai et al. 2024; Siletti et al. 2023). These resources can serve as training data for supervised classification models, allowing researchers to move beyond traditional marker gene matching toward automated label transfer. *CellTypist* is a prominent tool in this domain, offering logistic regression models trained on diverse tissues (Xu et al. 2023; Domínguez Conde et al. 2022). *CellTypist* is integrated within *scprocess* to provide a reproducible and efficient complementary approach to manual annotation. Additionally, *scprocess* provides an XGBoost model trained on adult human whole-brain dataset (Siletti et al. 2023), facilitating automated annotation of broad central nervous system cell types.

### Processing multiplexed samples

Multiplexing strategies are commonly used to scale up single cell experiments, allowing for simultaneous analysis of multiple samples within a single reaction. Common physical labeling approaches include tagging cells or nuclei in individual samples with lipid-bound or antibody-conjugated oligonucleotides (hashtag oligos, or HTOs) prior to pooling. Alternatively, in studies involving genetically distinct individuals, sample identities can be recovered based on natural genetic variation. *scprocess* supports the analysis of multiplexed samples by leveraging *simpleaf* (He and Patro 2023) to generate HTO count matrices from associated cDNA libraries. These results are then processed via the *Seurat HTODemux* function (Stoeckius et al. 2018) to assign cells to their samples of origin and identify technical multiplets. *scprocess* also accommodates outputs from external demultiplexing tools, enabling seamless processing of samples regardless of the multiplexing strategy employed.

### Subclustering

*scprocess* offers a subclustering feature that enables users to perform a second round of analysis on targeted cell populations. These subsets can be defined based on initial unsupervised clustering results, user-provided annotations or automated cell type labels. For selected cell subsets, *scprocess* generates pseudobulk profiles to reassess ambient contamination, identifies a new set of highly variable genes, and performs a secondary round of data integration and marker gene detection. This functionality is particularly valuable for resolving biological complexity that may be masked at the global scale. While initial clustering often captures broad cell lineages, subclustering can uncover subtle cellular heterogeneity such as transitory developmental stages or cell type activation states.

### Outputs

*scprocess* is designed to be agnostic to experimental design and therefore, does not include specialized analyses e.g. RNA velocity analysis or trajectory inference for time-series data. It preserves intermediate outputs at every major step and produces a collection of H5AD files containing filtered and integrated data, which can optionally be converted to *SingleCellExperiment* objects and used for a variety of downstream tasks. Additionally, *scprocess* generates multiple HTML reports from user-accessible RMarkdown files which can be easily customized to include additional diagnostic metrics tailored to the unique requirements of a specific dataset.

## Acknowledgements

We would like to thank Karsten Bach, Matthew Leipner, Atreya Choudhury, Pierre-Luc Germain and Miriam Pauline Busch, for their assistance in testing the pipeline and their constructive feedback. Additionally, we thank Caspar-Elias Glock for his advice on refining the doublet identification strategy.

## CRediT statement

MK: Methodology, Software development and testing, Data Curation, Writing – Original Draft, Writing – Review & Editing, Visualization, Project administration. JP: Methodology, Software development and testing. EB: Software development and testing. DG: Software development and testing. MR: Supervision, Writing – Review & Editing. WM: Conceptualization, Methodology, Software development and testing, Writing – Review & Editing, Visualization, Supervision, Funding acquisition.

