## Supplementary Table 1 for "scprocess: a pipeline for processing, integrating and visualising atlas-scale single cell data"

Data in this table was up-to-date on **2026-03-04**.

★ **Uses GPU**

| Method | scprocess | scRNAsequest | scRNAbox | SWANS | nf-core/scrnaseq | nf-core/scdownstream | Cellsnake |
|---|---|---|---|---|---|---|---|
| **Version** | v0.1.0 | / | v0.1.52.50 | v2.1.0 | 4.1.0 | / | v0.2.0.11 |
| **Paper** | this document | paper | paper | paper | / | / | paper |
| **Repo** | repo | repo | repo | repo | repo | repo | repo |
| **Docs** | docs | docs | docs | / | docs | docs | docs |
| **Implementation** | | | | | | | |
| **Workflow management system** | ✔ Snakemake | ✘ | ✘ | ✔ Snakemake | ✔ Nextflow | ✔ Nextflow | ✔ Snakemake |
| **Sequencing technology support** | 10x | 10x | 10x | droplet-based | 10x | droplet-based | droplet-based |
| **Input data** | FASTQ | HDF5, MEX | FASTQ | FASTQ, MEX | FASTQ | H5AD, SingleCellExperiment, Seurat, CSV | HDF5 |
| **Preprocessing features** | | | | | | | |
| **Mapping/UMI counting** | ✔ simpleaf | ✘ | ✔ CellRanger | ✔ CellRanger | ✔ CellRanger<br>Salmon-Alevin<br>STARsolo<br>Kallisto/Bustools | ✘ | ✘ |
| **Ambient RNA removal** | ✔ CellBender★<br>decontX | ✔ Cellbender★ | ✔ SoupX | ✔ SoupX | ✔ Cellbender★ | ✔ SoupX<br>CellBender★<br>decontX<br>scAR★ | ✘ |
| **Cell calling** | ✔ emptyDrops<br>Cellbender★<br>barcodeRanks | ✘ | ✔ CellRanger | ✔ CellRanger | ✔ Cellbender★<br>CellRanger<br>STARsolo<br>Kallisto/Bustools | ✔ Cellbender★ | ✘ |
| **Doublet detection** | ✔ scDblFinder<br>HTOdemux<br>removal of doublet-enriched clusters | ✔ scDblFinder | ✔ DoubletFinder<br>Seurat-MULTIseqDemux | ✔ DoubletFinder | ✘ | ✔ SOLO ★<br>scrublet<br>DoubletDetection<br>SCDS | ✔ DoubletFinder |
| **QC thresholds** | ✔ user defined | ✔ user defined | ✔ user defined | ✔ user defined | ✘ | ✔ user defined | ✔ user defined<br>MAD<br>miQC |
| **QC filtering** | ✔ library size<br>detected features<br>mito %<br>spliced % | ✔ library size<br>detected features<br>mito % | ✔ library size<br>detected features<br>mito %<br>ribosomal % | ✔ library size<br>detected features<br>mito %<br>ribosomal % | ✘ | ✔ library size<br>detected features<br>mito % | ✔ library size<br>detected features<br>mito % |
| **HVG selection** | ✔ Seurat-based (includes per-batch option; optimized for handling large datasets)<br>can exclude ambient genes<br>can exclude specified genes | ✔ Seurat::SelectIntegrationFeatures (includes per-batch option) | ✔ Seurat::SelectIntegrationFeatures (includes per-batch option) | ✔ Seurat::FindVariableFeatures (whole dataset only) | ✘ | ✔ scanpy.pp.highly_variable_genes<br>Seurat::SelectIntegrationFeatures (includes per-batch option)<br>can exclude specified genes | ✔ Seurat::FindVariableFeatures (whole dataset only) |
| **Normalization** | ✔ Log normalization | ✔ Log normalization<br>SCTransform | ✔ Log normalization | ✔ Log normalization<br>SCTransform | ✘ | ✔ Log normalization<br>SCTransform | ✔ Log normalization |
| **UMAP** | ✔ Scanpy<br>RAPIDS-singlecell★ | ✔ Seurat<br>Scanpy | ✔ Seurat | ✔ Seurat | ✘ | ✔ Scanpy | ✔ Seurat |
| **Batch correction** | ✔ Harmony (RAPIDS-singlecell)★<br>Harmony (Scanpy) | ✔ Seurat-CCA/RPCA<br>Liger<br>Harmony (R) | ✔ Seurat-CCA | ✔ Seurat-CCA/RPCA<br>Harmony (R) | ✘ | ✔ scVI★<br>scANVI★<br>Harmony (Scanpy)<br>Combat (Scanpy)<br>Seurat-CCA<br>BBKNN | ✔ Seurat-CCA/RPCA |
| **Clustering** | ✔ Leiden (RAPIDS-singlecell)★<br>Leiden (Scanpy) | ✔ Louvain (Seurat) | ✔ Louvain (Seurat) | ✔ Louvain (Seurat | ✘ | ✔ Leiden (Scanpy) | ✔ Louvain (Seurat) |
| **Marker gene detection** | ✔ edgeR (pseudobulk) | ✘ | ✔ Seurat::FindAllMarkers (cell level) | ✔ Seurat::FindConservedMarkers (cell level) | ✘ | ✔ scanpy.tl.rank_genes_groups (cell level) | ✔ Seurat::FindAllMarkers (cell level) |
| **Marker genes pathway analysis** | ✔ fgsea | ✘ | ✔ enrichR | ✘ | ✘ | ✘ | ✔ clusterProfiler |
| **Cell type annotation** | ✔ CellTypist<br>XGBoost (human brain only) | ✔ Seurat::TransferData<br>scRef | ✔ Seurat::TransferData | ✔ Seurat::TransferData<br>Azimuth | ✘ | ✔ CellTypist<br>SingleR | ✔ CellTypist<br>SingleR |
| **Subclustering** | ✔ | ✔ | ✘ | ✘ | ✘ | ✔ | ✘ |
| **Sample demultiplexing** | ✔ Seurat::HTODemux<br>user defined cell-donor assignments | ✘ | ✔ Seurat::MULTIseqDemux | ✘ | ✘ | ✘ | ✘ |
| **Additional features** | | | | | | | |
| | | Differential gene expression analysis | Diffferential gene expression analysis (cell-based using Seurat::FindMarkers and MAST, sample-based using DESeq2) | Option to run multiple different combinations of methods at once, differential gene expression analysis + GSEA (fgsea) on DEGs, trajectory analysis using Monocle3, | | | Metagenome analysis using Kraken2, cell-cell communication analysis using CellChat |
