## Supplementary File 1 for "scprocess: a pipeline for processing, integrating and visualising atlas-scale single cell data"

### Selected scprocess outputs for Pineda et al. 2024

### Introduction

To demonstrate the scalability and performance of `scprocess` on a large-scale single cell dataset, we processed 149 samples from the following study:

*Pineda, S. Sebastian, Hyeseung Lee, Maria J. Ulloa-Navas, et al. 2024. "Single-Cell Dissection of the Human Motor and Prefrontal Cortices in ALS and FTLD." Cell 187 (8): 1971–1989.e16.*

FASTQ files were downloaded from Gene Expression Omnibus (GSE174332) and used as input to `scprocess`. All core steps of the pipeline were executed, as well as cell type labeling and subclustering of myeloid cells and neurons, using the following configuration file:

```yaml
project:
  proj_dir:         /pstore/data/brain-sc-analysis/studies/pineda_2024
  arv_uuids:        [not shown]
  arv_instance:     arkau
  show_arv_uuids:   False
  full_tag:         pineda_2024
  short_tag:        pineda
  your_name:        Will Macnair
  affiliation:      Neuroscience and Rare Diseases, Roche Innovation Center, Basel, Switzerland
  sample_metadata:  data/metadata/pineda_2024_harmonized_metadata.csv
  metadata_vars:    [diagnosis, diagnosis_subtype, original_tissue_annotation]
  ref_txome:        human_2024
  date_stamp:       "2026-02-28"
  custom_sample_params: data/metadata/custom_params_pineda_2024.yaml
ambient:
  ambient_method:   cellbender
qc:
  qc_max_mito:      0.10
  qc_max_splice:    0.75
marker_genes:
  mkr_custom_genesets:
  - name: human_brain
label_celltypes:
  - labeller:       scprocess
    model:          human_cns
zoom:
  - data/metadata/zoom_params_pineda_2024_myeloid.yaml
  - data/metadata/zoom_params_pineda_2024_neurons.yaml
resources:
  retries:                          1
  gb_render_html_label_celltypes:   32
  gb_render_html_zoom:              32
  gb_run_integration:               64
  gb_run_marker_genes:              98
  gb_zoom_run_marker_genes:         98
  mins_render_html_label_celltypes: 60
  mins_run_cellbender:              180
  mins_run_mapping:                 120
  mins_run_marker_genes:            180
  mins_zoom_run_marker_genes:       120
```

While `scprocess` automatically generates comprehensive HTML reports for each pipeline stage, the relevant diagnostic plots have been consolidated into this document for brevity. Note that where the original interactive reports use tabs to display e.g. sample-specific or resolution-specific plots, we provide a single representative example here. The complete, interactive report is available at https://marusakod.github.io/scprocess__pineda_2024/.

### Read mapping diagnostics

#### Barcode rank plots

For each sample, barcodes are ranked in descending order based on their library size (number of unique molecular identifiers (UMIs)). Typically, one can observe two plateaus in the barcode-rank curve: the first corresponds to droplets containing cells with high RNA content, while the second represents empty droplets containing ambient RNA. Key transition points on the curve (shin and knee points), are annotated with horizontal lines. Two main parameters inferred by `scprocess` based on transition points are `expected_cells` and `empty_plateau_middle`.`empty_plateau_middle` should extend a few thousand barcodes into the second plateau. The absence of a clear empty plateau indicates poor sample quality due to ambient RNA contamination.

##### Ordered by slope ratio

The slope ratio helps identify low-quality samples with high ambient RNA contamination. This ratio is calculated by dividing the slope of the barcode-rank curve in the empty droplet plateau by the slope at the first inflection point. Samples are ordered from highest to lowest based on the slope ratio.

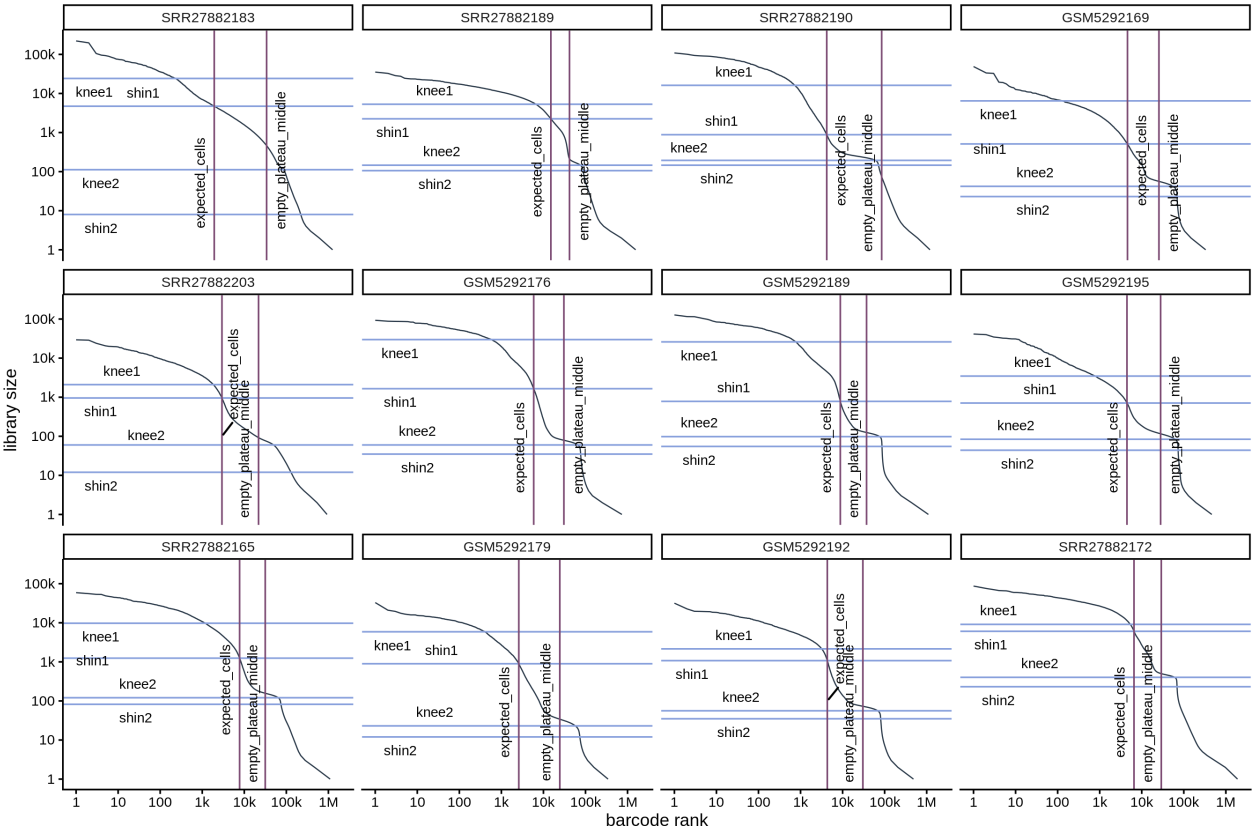

##### Dotplot with diagnotic ratios

Both the slope ratio and the expected cells to empty droplets ratio are displayed in a dot plot, where each dot represents a sample. Outliers are labeled with their corresponding sample_id values.

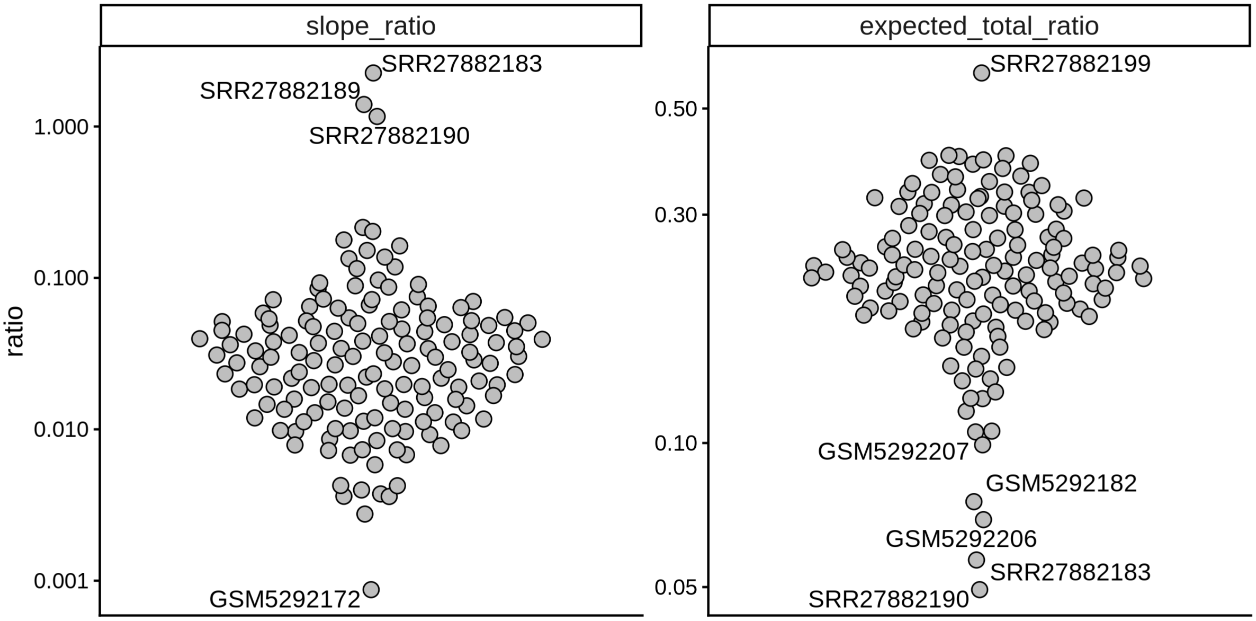

#### Library sizes and splice read proportions: comparison between cells and empty droplets across samples

This plot shows the distribution of UMIs and spliced read proportions for both empty droplets and expected cells, split by sample. The UMI counts in empty droplets serve as an indicator of ambient RNA contamination in the dataset.

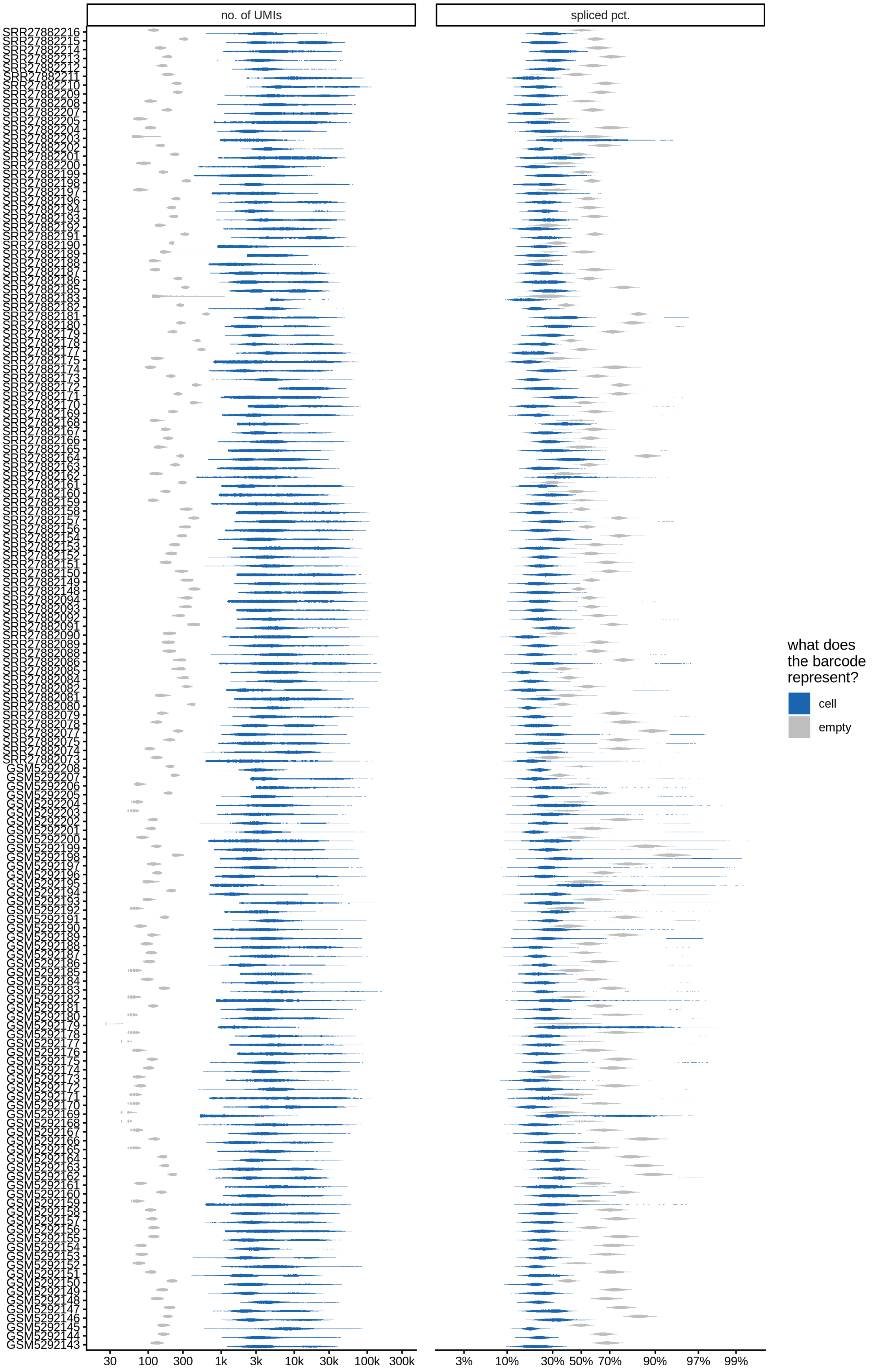

no. of UMIs
spliced pct.
30
100
300
1k
3k
10k
30k
100k
300k
3%
10%
30%
50%
70%
90%
97%
99%
what does
the barcode
represent?
cell
empty

### Ambient RNA removal diagnostics

#### Distributions of spliced pct. in empty and cell barcodes

The plots show the relationship between the number of UMIs and the percentage of spliced reads, separated by sample, both before and after ambient RNA removal. Ambient RNA is typically characterized by a high proportion of spliced reads, making the spliced percentage a useful metric for evaluating the effectiveness of an ambient RNA removal method. By examining changes in spliced percentages, we can assess how well the method performed in reducing ambient RNA contamination.

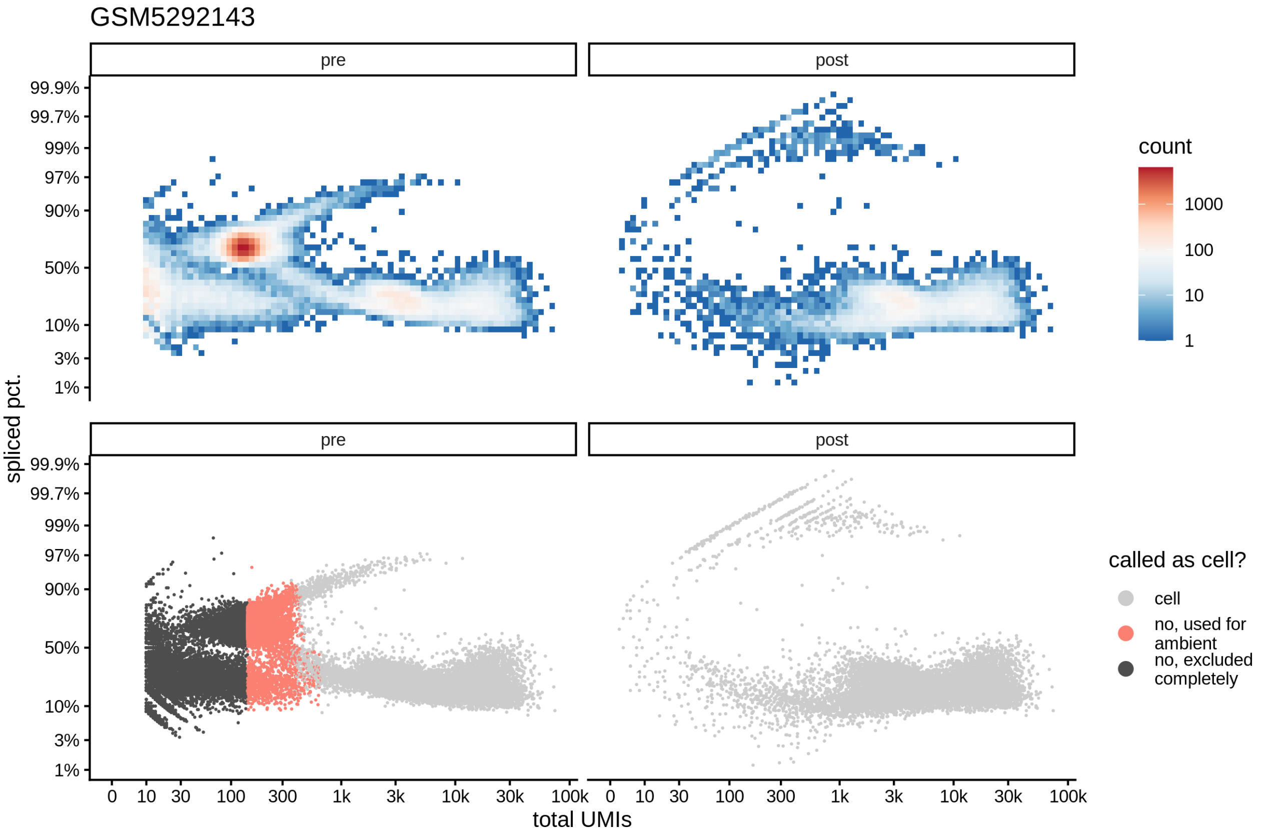

#### How many reads were removed as ambient?

Plots show what proportion of reads were removed from all barcodes called as cells.

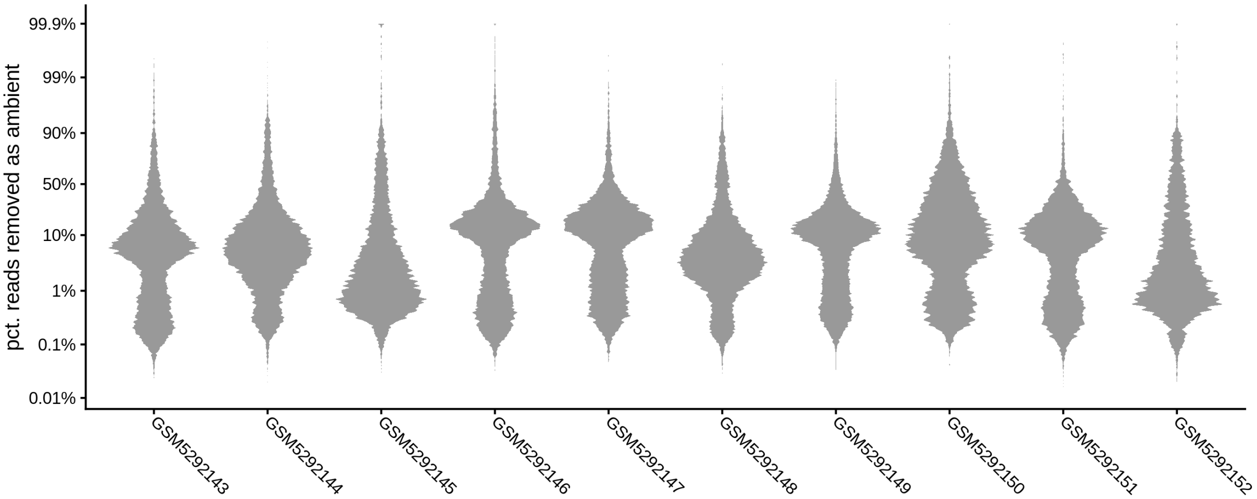

#### Samples excluded by ambient removal step

CellBender includes all barcodes in the analysis up to the `total_droplets` threshold covering both cell-containing droplets and empty droplets. If CellBender calls the majority of these included droplets as cells, it may indicate an underlying issue. This typically occurs in low-quality samples where cell-containing barcodes and empty droplets cannot be clearly distinguished in the barcode rank plot. The table below shows the proportion of included droplets that were classified as cells by CellBender. Samples where this proportion exceeded % were excluded from further analysis.

| sample_id | total_droplets | kept_droplets | pct_kept | bad_run |
|---|---:|---:|---:|---|
| GSM5292143 | 35173 | 11679 | 33.2 | FALSE |
| GSM5292144 | 37914 | 16306 | 43.0 | FALSE |
| GSM5292145 | 29983 | 7186 | 24.0 | FALSE |
| GSM5292146 | 35361 | 12222 | 34.6 | FALSE |
| GSM5292147 | 37893 | 13946 | 36.8 | FALSE |
| GSM5292148 | 36487 | 12769 | 35.0 | FALSE |
| GSM5292149 | 41396 | 16483 | 39.8 | FALSE |
| GSM5292150 | 40048 | 14190 | 35.4 | FALSE |
| GSM5292151 | 37819 | 11602 | 30.7 | FALSE |
| GSM5292152 | 33970 | 5836 | 17.2 | FALSE |
| GSM5292153 | 35151 | 8837 | 25.1 | FALSE |
| GSM5292154 | 35114 | 7581 | 21.6 | FALSE |
| GSM5292155 | 36153 | 7526 | 20.8 | FALSE |
| GSM5292156 | 34555 | 9000 | 26.0 | FALSE |
| GSM5292157 | 40829 | 9877 | 24.2 | FALSE |
| GSM5292158 | 37305 | 11266 | 30.2 | FALSE |
| GSM5292159 | 28882 | 8232 | 28.5 | FALSE |
| GSM5292160 | 40863 | 15402 | 37.7 | FALSE |
| GSM5292161 | 30376 | 9463 | 31.2 | FALSE |
| GSM5292162 | 33760 | 11168 | 33.1 | FALSE |
| GSM5292163 | 38375 | 15452 | 40.3 | FALSE |
| GSM5292164 | 38757 | 13156 | 33.9 | FALSE |
| GSM5292165 | 35933 | 14462 | 40.2 | FALSE |
| GSM5292166 | 31982 | 10335 | 32.3 | FALSE |
| GSM5292167 | 29697 | 8779 | 29.6 | FALSE |
| GSM5292168 | 27530 | 4241 | 15.4 | FALSE |
| GSM5292169 | 25636 | 10110 | 39.4 | FALSE |

| sample_id | total_droplets | kept_droplets | pct_kept | bad_run |
|---|---|---|---|---|
| GSM5292170 | 28106 | 4603 | 16.4 | FALSE |
| GSM5292171 | 26013 | 3370 | 13.0 | FALSE |
| GSM5292172 | 31901 | 5743 | 18.0 | FALSE |
| GSM5292173 | 27815 | 5683 | 20.4 | FALSE |
| GSM5292174 | 32868 | 8040 | 24.5 | FALSE |
| GSM5292175 | 35862 | 8644 | 24.1 | FALSE |
| GSM5292176 | 30626 | 9818 | 32.1 | FALSE |
| GSM5292177 | 27604 | 5668 | 20.5 | FALSE |
| GSM5292178 | 34758 | 7913 | 22.8 | FALSE |
| GSM5292179 | 24487 | 7138 | 29.2 | FALSE |
| GSM5292180 | 35542 | 8810 | 24.8 | FALSE |
| GSM5292181 | 33774 | 7517 | 22.3 | FALSE |
| GSM5292182 | 27916 | 3716 | 13.3 | FALSE |
| GSM5292183 | 13008 | 1856 | 14.3 | FALSE |
| GSM5292184 | 31384 | 6391 | 20.4 | FALSE |
| GSM5292185 | 28720 | 6241 | 21.7 | FALSE |
| GSM5292186 | 36164 | 9651 | 26.7 | FALSE |
| GSM5292187 | 33555 | 8364 | 24.9 | FALSE |
| GSM5292188 | 34637 | 6880 | 19.9 | FALSE |
| GSM5292189 | 37125 | 14088 | 37.9 | FALSE |
| GSM5292190 | 31745 | 7506 | 23.6 | FALSE |
| GSM5292191 | 37785 | 9860 | 26.1 | FALSE |
| GSM5292192 | 30344 | 8945 | 29.5 | FALSE |
| GSM5292193 | 33076 | 8060 | 24.4 | FALSE |
| GSM5292194 | 39819 | 14566 | 36.6 | FALSE |
| GSM5292195 | 28079 | 6666 | 23.7 | FALSE |
| GSM5292196 | 35030 | 8912 | 25.4 | FALSE |
| GSM5292197 | 37197 | 11653 | 31.3 | FALSE |
| GSM5292198 | 26739 | 9722 | 36.4 | FALSE |
| GSM5292199 | 36340 | 9856 | 27.1 | FALSE |
| GSM5292200 | 33537 | 9522 | 28.4 | FALSE |
| GSM5292201 | 34889 | 7267 | 20.8 | FALSE |
| GSM5292202 | 34340 | 8527 | 24.8 | FALSE |
| GSM5292203 | 28656 | 6256 | 21.8 | FALSE |
| GSM5292204 | 28264 | 5792 | 20.5 | FALSE |
| GSM5292205 | 33974 | 8918 | 26.2 | FALSE |
| GSM5292206 | 24306 | 5001 | 20.6 | FALSE |
| GSM5292207 | 26218 | 4257 | 16.2 | FALSE |
| GSM5292208 | 35771 | 9686 | 27.1 | FALSE |
| SRR27882073 | 29280 | 8385 | 28.6 | FALSE |
| SRR27882074 | 33844 | 10155 | 30.0 | FALSE |
| SRR27882075 | 40876 | 18327 | 44.8 | FALSE |
| SRR27882077 | 39344 | 16662 | 42.3 | FALSE |
| SRR27882078 | 36400 | 11305 | 31.1 | FALSE |
| SRR27882079 | 36652 | 12050 | 32.9 | FALSE |
| SRR27882080 | 34560 | 12622 | 36.5 | FALSE |
| SRR27882081 | 33960 | 11481 | 33.8 | FALSE |
| SRR27882082 | 35856 | 14312 | 39.9 | FALSE |
| SRR27882084 | 39543 | 7860 | 19.9 | FALSE |
| SRR27882085 | 40989 | 8579 | 20.9 | FALSE |
| SRR27882086 | 41889 | 9570 | 22.8 | FALSE |
| SRR27882088 | 42788 | 9514 | 22.2 | FALSE |

| sample_id | total_droplets | kept_droplets | pct_kept | bad_run |
|---|---|---|---|---|
| SRR27882089 | 41816 | 8876 | 21.2 | FALSE |
| SRR27882090 | 41326 | 9341 | 22.6 | FALSE |
| SRR27882091 | 45593 | 13498 | 29.6 | FALSE |
| SRR27882092 | 44460 | 14009 | 31.5 | FALSE |
| SRR27882093 | 44478 | 13993 | 31.5 | FALSE |
| SRR27882094 | 45151 | 12893 | 28.6 | FALSE |
| SRR27882148 | 40254 | 8463 | 21.0 | FALSE |
| SRR27882149 | 41538 | 10412 | 25.1 | FALSE |
| SRR27882150 | 38367 | 10916 | 28.5 | FALSE |
| SRR27882151 | 40678 | 10083 | 24.8 | FALSE |
| SRR27882152 | 40305 | 9619 | 23.9 | FALSE |
| SRR27882153 | 37919 | 10088 | 26.6 | FALSE |
| SRR27882154 | 40897 | 12161 | 29.7 | FALSE |
| SRR27882156 | 40536 | 10797 | 26.6 | FALSE |
| SRR27882157 | 40451 | 10470 | 25.9 | FALSE |
| SRR27882158 | 37214 | 8806 | 23.7 | FALSE |
| SRR27882159 | 35887 | 9297 | 25.9 | FALSE |
| SRR27882160 | 35410 | 10892 | 30.8 | FALSE |
| SRR27882161 | 29457 | 7371 | 25.0 | FALSE |
| SRR27882162 | 6287 | 1703 | 27.1 | FALSE |
| SRR27882163 | 38995 | 14087 | 36.1 | FALSE |
| SRR27882164 | 44100 | 16831 | 38.2 | FALSE |
| SRR27882165 | 31510 | 12266 | 38.9 | FALSE |
| SRR27882166 | 31296 | 7292 | 23.3 | FALSE |
| SRR27882167 | 35618 | 10898 | 30.6 | FALSE |
| SRR27882168 | 32262 | 11076 | 34.3 | FALSE |
| SRR27882169 | 35378 | 14293 | 40.4 | FALSE |
| SRR27882170 | 29529 | 12479 | 42.3 | FALSE |
| SRR27882171 | 38752 | 15852 | 40.9 | FALSE |
| SRR27882172 | 29158 | 18812 | 64.5 | FALSE |
| SRR27882173 | 35731 | 11511 | 32.2 | FALSE |
| SRR27882174 | 38376 | 11363 | 29.6 | FALSE |
| SRR27882175 | 27856 | 6665 | 23.9 | FALSE |
| SRR27882177 | 34173 | 13445 | 39.3 | FALSE |
| SRR27882178 | 35427 | 14195 | 40.1 | FALSE |
| SRR27882179 | 34644 | 11646 | 33.6 | FALSE |
| SRR27882180 | 41815 | 19204 | 45.9 | FALSE |
| SRR27882181 | 38825 | 16311 | 42.0 | FALSE |
| SRR27882182 | 38542 | 13042 | 33.8 | FALSE |
| SRR27882183 | 33905 | 22026 | 65.0 | FALSE |
| SRR27882185 | 42864 | 15173 | 35.4 | FALSE |
| SRR27882186 | 37651 | 9973 | 26.5 | FALSE |
| SRR27882187 | 40324 | 10414 | 25.8 | FALSE |
| SRR27882188 | 34575 | 8310 | 24.0 | FALSE |
| SRR27882189 | 41612 | 37137 | 89.2 | FALSE |
| SRR27882190 | 85000 | 69389 | 81.6 | FALSE |
| SRR27882191 | 27170 | 6038 | 22.2 | FALSE |
| SRR27882192 | 30063 | 6396 | 21.3 | FALSE |
| SRR27882193 | 30288 | 6841 | 22.6 | FALSE |
| SRR27882194 | 30397 | 5340 | 17.6 | FALSE |
| SRR27882196 | 27661 | 5464 | 19.8 | FALSE |
| SRR27882197 | 27852 | 5832 | 20.9 | FALSE |

| sample_id | total_droplets | kept_droplets | pct_kept | bad_run |
|---|---|---|---|---|
| SRR27882198 | 27794 | 5476 | 19.7 | FALSE |
| SRR27882199 | 47047 | 28539 | 60.7 | FALSE |
| SRR27882200 | 41117 | 11334 | 27.6 | FALSE |
| SRR27882201 | 40758 | 14323 | 35.1 | FALSE |
| SRR27882202 | 33113 | 8251 | 24.9 | FALSE |
| SRR27882203 | 21813 | 5242 | 24.0 | FALSE |
| SRR27882204 | 41305 | 15855 | 38.4 | FALSE |
| SRR27882205 | 42521 | 10151 | 23.9 | FALSE |
| SRR27882207 | 33634 | 8062 | 24.0 | FALSE |
| SRR27882208 | 40549 | 10146 | 25.0 | FALSE |
| SRR27882209 | 32569 | 9768 | 30.0 | FALSE |
| SRR27882210 | 32448 | 6535 | 20.1 | FALSE |
| SRR27882211 | 36596 | 9993 | 27.3 | FALSE |
| SRR27882212 | 39104 | 10227 | 26.2 | FALSE |
| SRR27882213 | 36336 | 9662 | 26.6 | FALSE |
| SRR27882214 | 32912 | 8011 | 24.3 | FALSE |
| SRR27882215 | 34203 | 7496 | 21.9 | FALSE |
| SRR27882216 | 40307 | 10338 | 25.6 | FALSE |

### Cell-level QC

#### Sample QC metrics

An overview of quality control (QC) metrics for all samples, both before and after filtering. The QC metrics include:

- `no. of cells` : Total number of barcodes called as cells for each sample.
- `no. of UMIs` : Sum of counts across all features for each cell with log-transformed values displayed.
- `no. of genes` : Number of unique features with non-zero counts for each cell with log-transformed values displayed.
- `mito pct.` : The proportion of reads that mapped to genes in the mitochondrial genome. High proportions are indicative of poor-quality cells (compromised membranes allow individual RNA molecules to escape while retaining mitochondria, leading to an increased relative abundance of mitochondrial transcript) or nuclei (incomplete removal of cytoplasm). Logit-transformed values are displayed as applying the logit (inverse logistic) transformation to mitochondrial proportions yields approximately Gaussian distributions.
- `spliced pct.` : The proportion of spliced reads. In single-nuclei RNA sequencing, high spliced proportions may indicate inadequate removal of cytoplasmic material from the nuclei. Logit-transformed values are displayed as applying the logit (inverse logistic) transformation to spliced proportions results in approximately Gaussian distributions.

**Pre-QC**

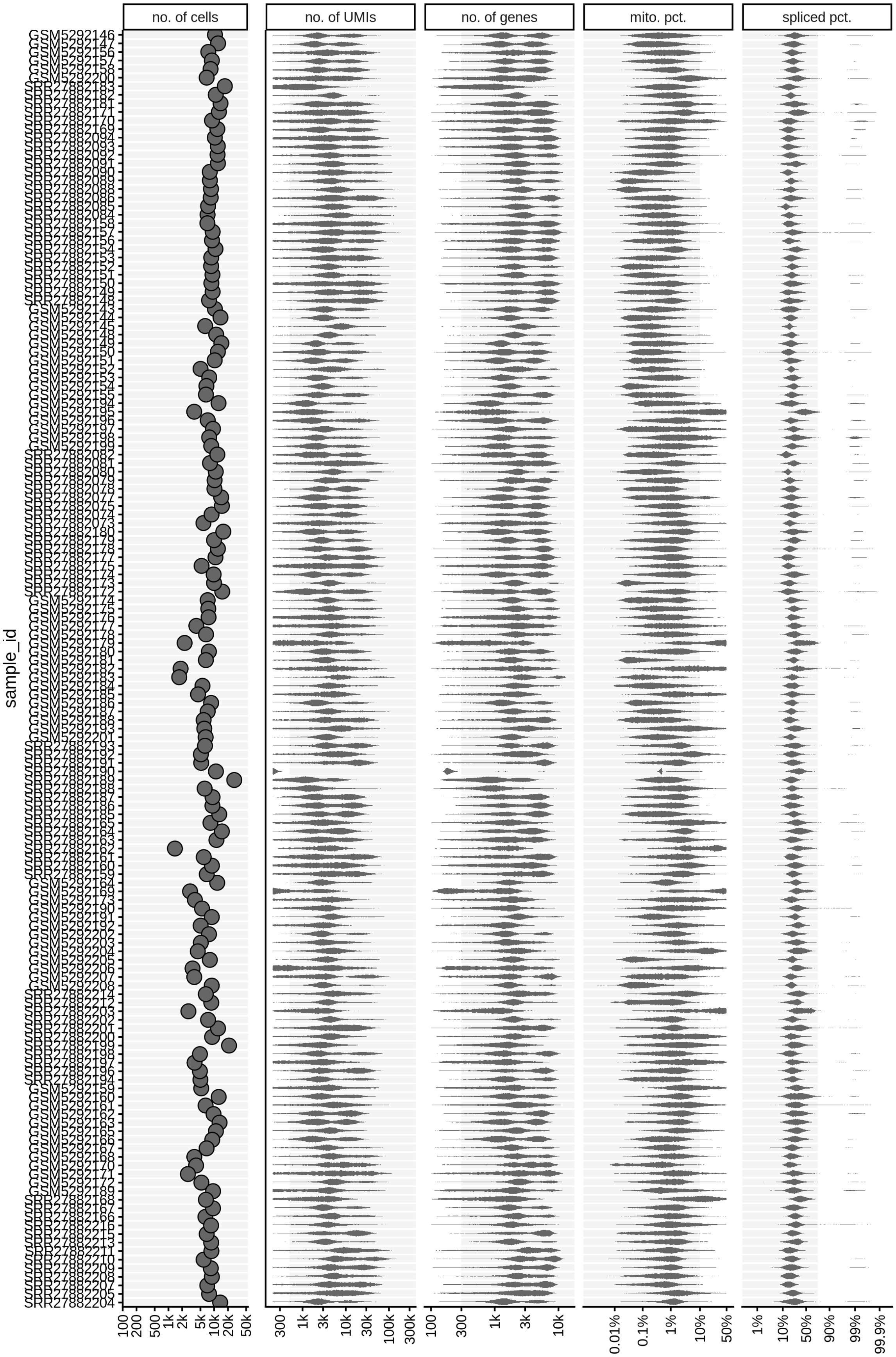
no. of cells
no. of UMIs
no. of genes
mito. pct.
spliced pct.
sample_id

**After light QC**

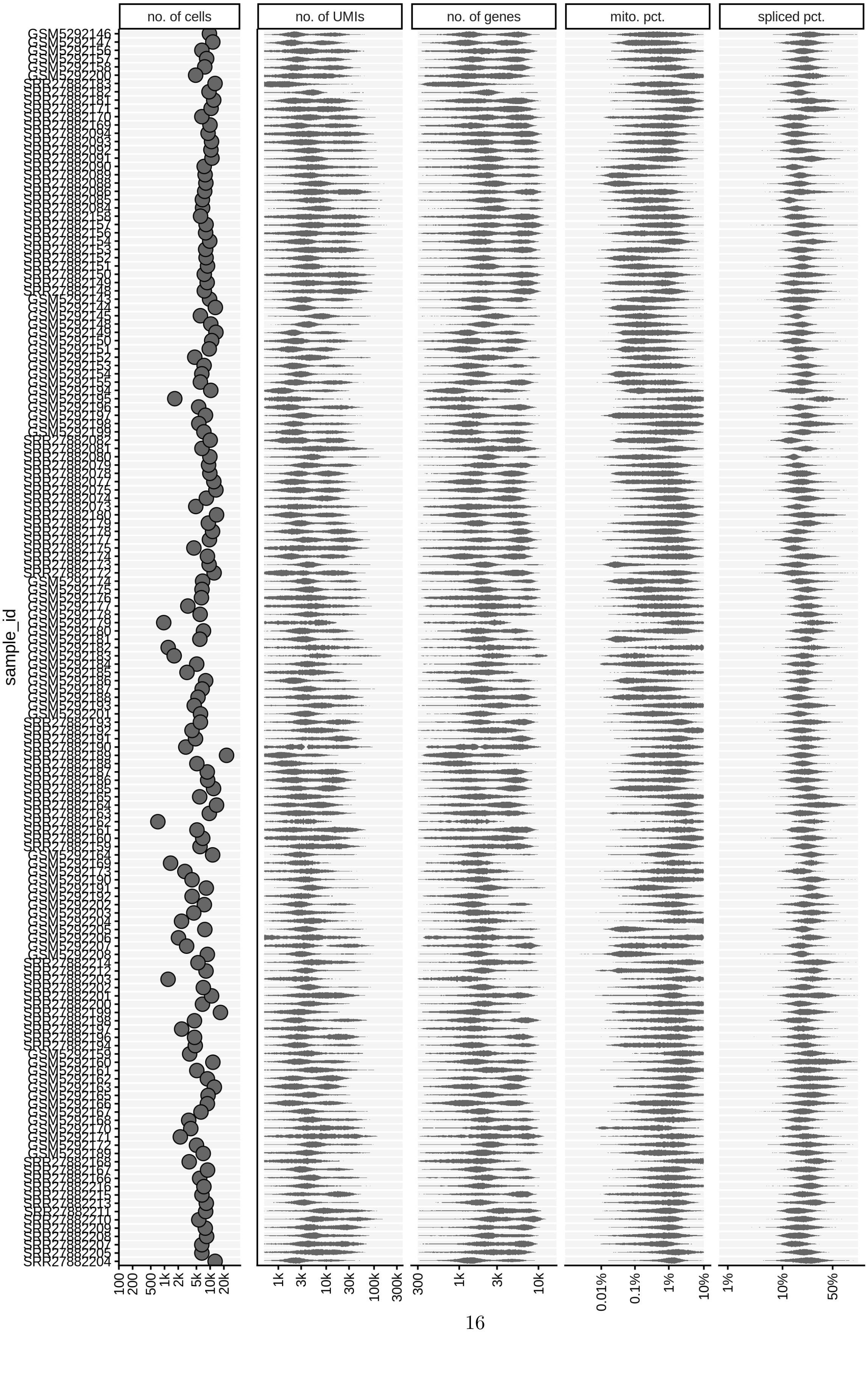

#### QC metrics scatterplots

Pairwise relationships of QC metrics at individual cell level.

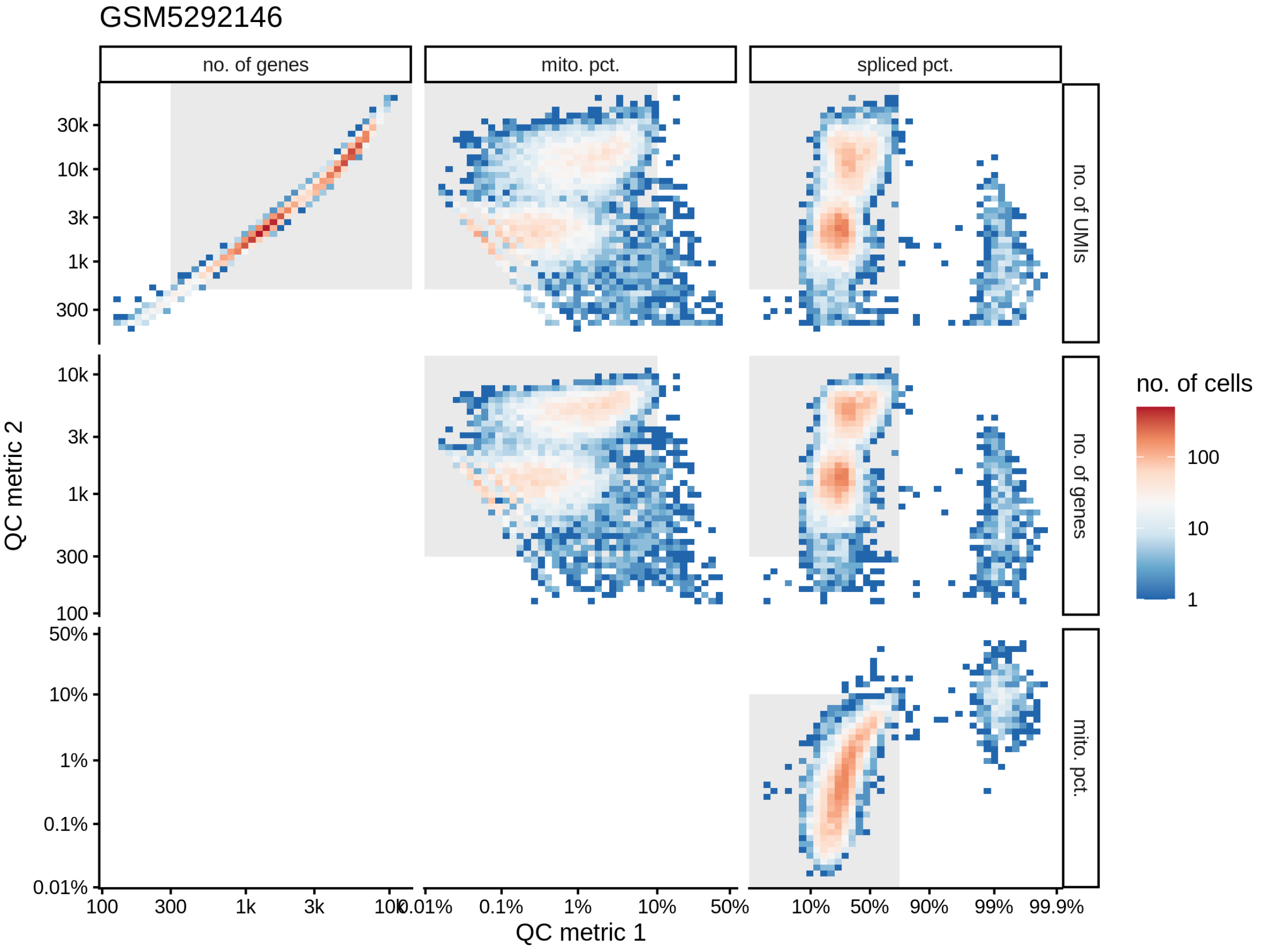

#### What were the reasons for excluding barcodes

An upset plot is generated for each sample to visualize the reasons for barcode exclusion. The plot shows the overlap between different exclusion criteria, with vertical bars representing the size of each intersection. Plots are omitted for samples where no barcodes were excluded.

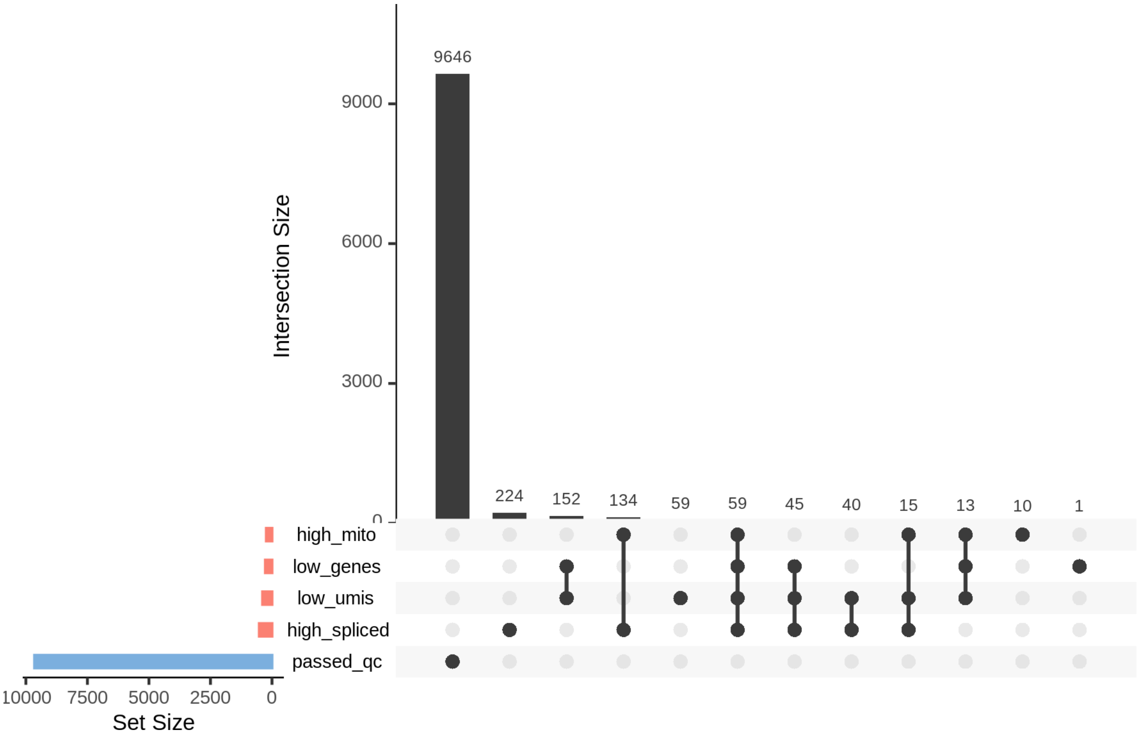

Intersection Size
9646
224
152
134
59
59
45
40
15
13
10
1
9000
6000
3000
0
high_mito
low_genes
low_umis
high_spliced
passed_qc
10000 7500 5000 2500 0
Set Size

### HVG selection diagnostics

The selection of highly variable genes is a crucial step in the integration process and in identifying clusters in single-cell experiments. However, contamination from ambient RNA can act as a significant confounding factor. This report provides plots to evaluate the influence of ambient RNA on the selection of highly variable genes and its potential impact on downstream analyses.

#### Are highly variable genes "ambient"?

The plot shows the extent to which highly variable genes identified using the `Seurat` VST method may be influenced by ambient RNA. The x-axis represents the `log2fc` values derived from the ambient gene estimation step in `scprocess` (see `scprocess` documentation for more details), while the y-axis shows the trend-normalized variance calculated using the `Seurat` VST method. Each point represents a gene, annotated based on whether it is among the top HVGs and whether it is identified as "ambient" by the ambient gene detection step. Labelled genes are top 20 with highest mean variance, including genes that were not included as HVGs because of high expression in empty droplets.

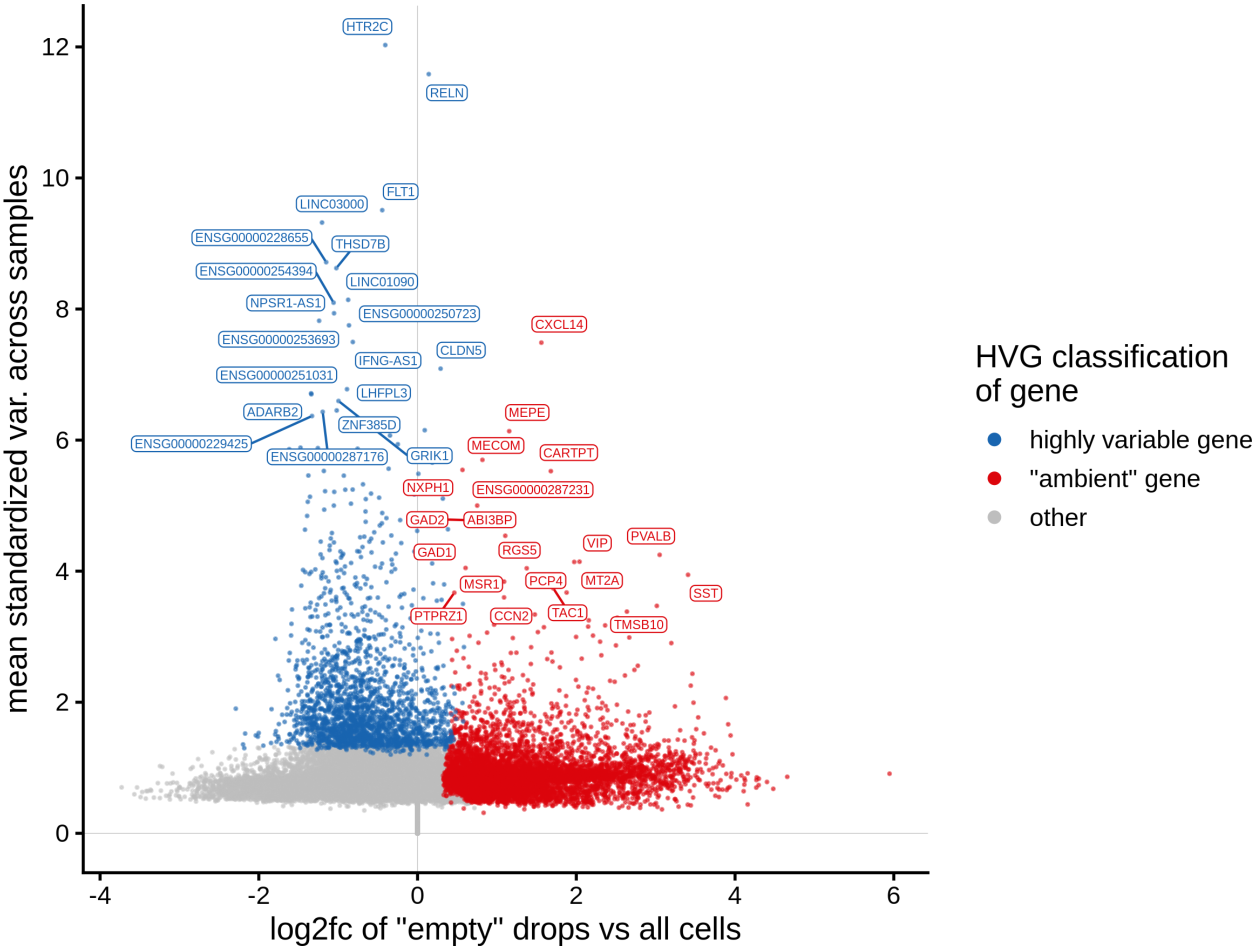

#### Which genes are "ambient"?

The plot shows the results of the `scprocess` ambient gene detection procedure. The y-axis is the `-log10` nominal p-value, with a dotted line indicating the threshold where the adjusted p-value is sufficiently small ($< 0.01$). Multiple plots are shown, each corresponding to a different minimum expression level filter applied to the ambient profiles.

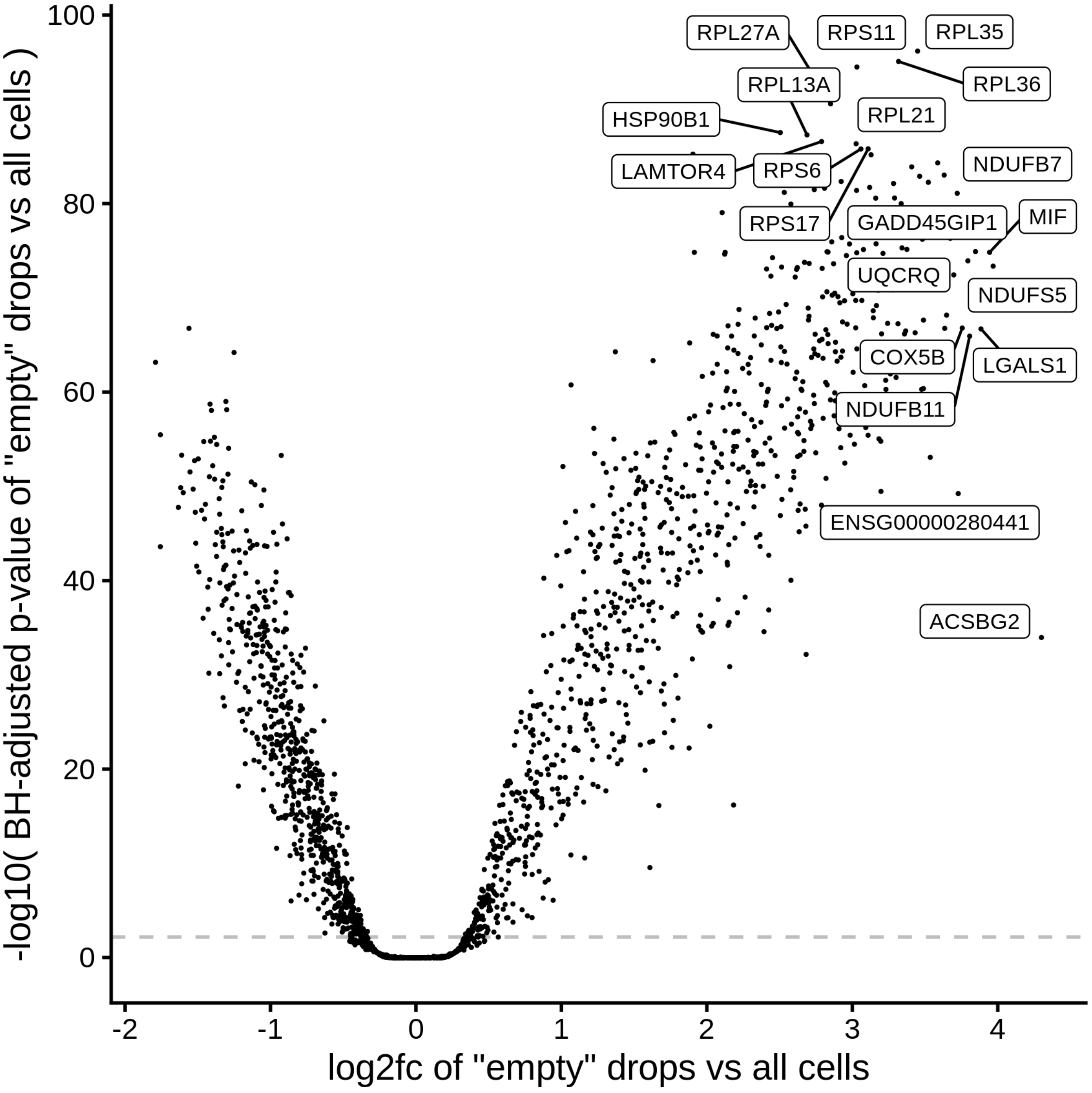

##### Which genes are variable across the ambient profiles?

Examining how ambient genes vary across different samples can provide valuable insights. Such variation may highlight cases where certain samples require distinct treatment, for example, if case and control samples consistently exhibit different ambient profiles.

The heatmaps display the pseudobulk expression of various genes across the ambient profiles of each sample in the dataset. The genes shown are the top 40 genes with the highest variance across ambient profiles:

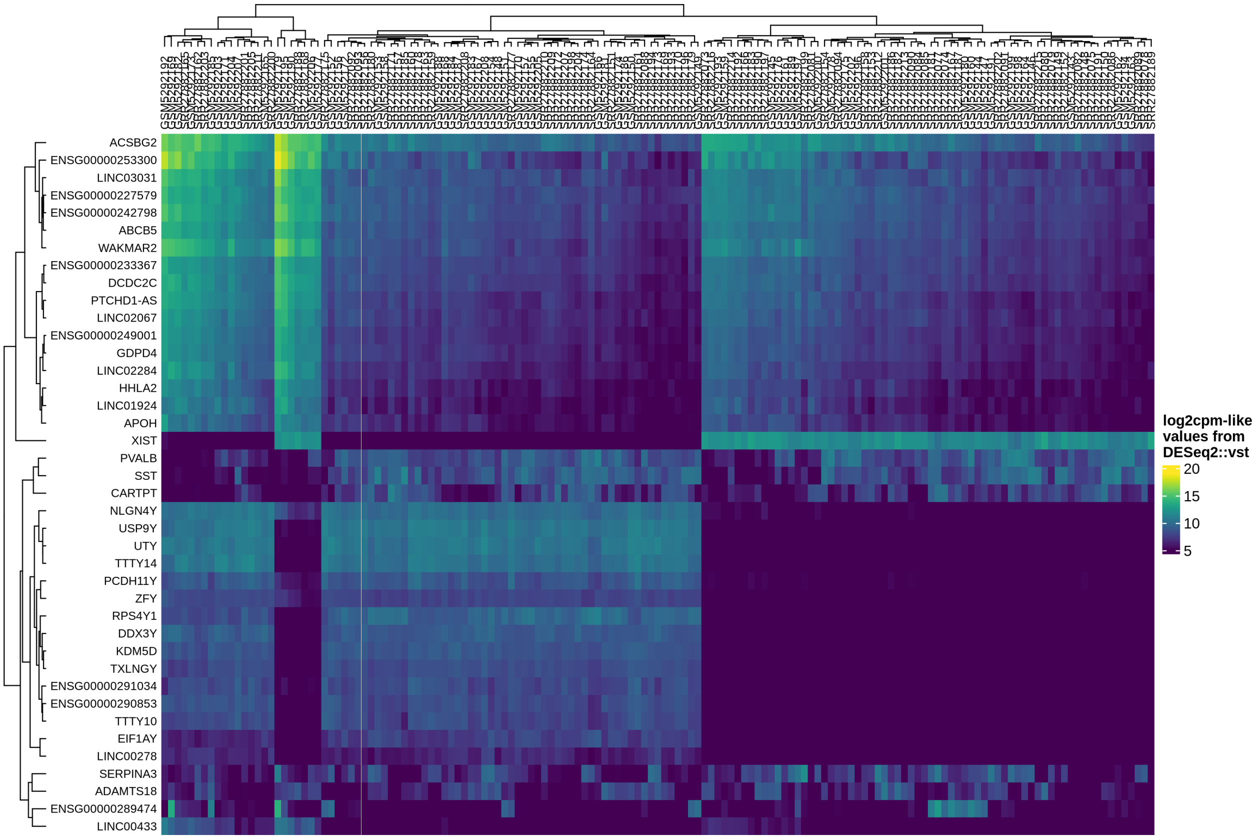
ACSBG2
ENSG00000253300
LINC03031
ENSG00000227579
ENSG00000242798
ABCB5
WAKMAR2
ENSG00000233367
DCDC2C
PTCHD1-AS
LINC02067
ENSG00000249001
GDPD4
LINC02284
HHLA2
LINC01924
APOH
XIST
PVALB
SST
CARTPT
NLGN4Y
USP9Y
UTY
TTTY14
PCDH11Y
ZFY
RPS4Y1
DDX3Y
KDM5D
TXLNGY
ENSG00000291034
ENSG00000290853
TTTY10
EIF1AY
LINC00278
SERPINA3
ADAMTS18
ENSG00000289474
LINC00433
log2cpm-like values from DESeq2::vst
20
15
10
5

### Integration diagnostics

Cells that passed quality control filtering were integrated with `Harmony` together with doublets identified using `scDblfinder`. Clustering of cells at a high resolution is then performed. Cells that appear in clusters which are enriched in doublets are excluded from further analysis. Integration of cells and clustering is repeated after removing doublet enriched clusters.

#### Doublets over UMAP

The plot displays a binned UMAP with the proportion of doublets as well as the number of doublets in each bin.

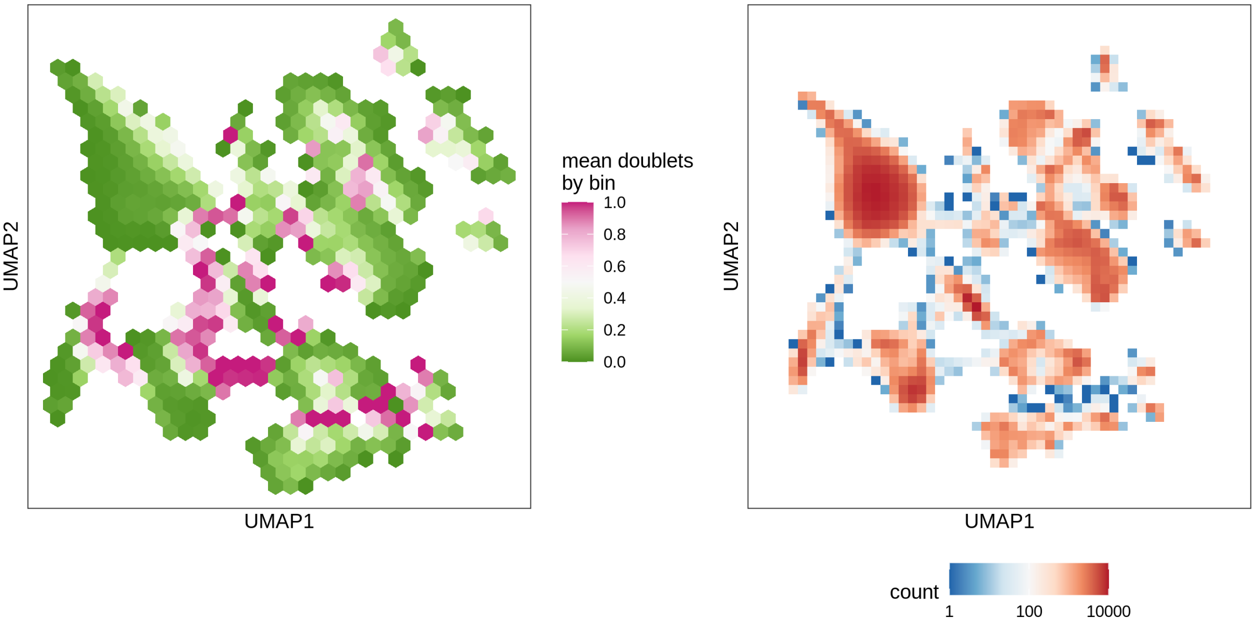

#### Doublet proportions in clusters

The plot shows the proportion of doublets for each cluster in relation to the total number of cells in that cluster. Clusters with a doublet proportion exceeding 50% are excluded from further analysis.

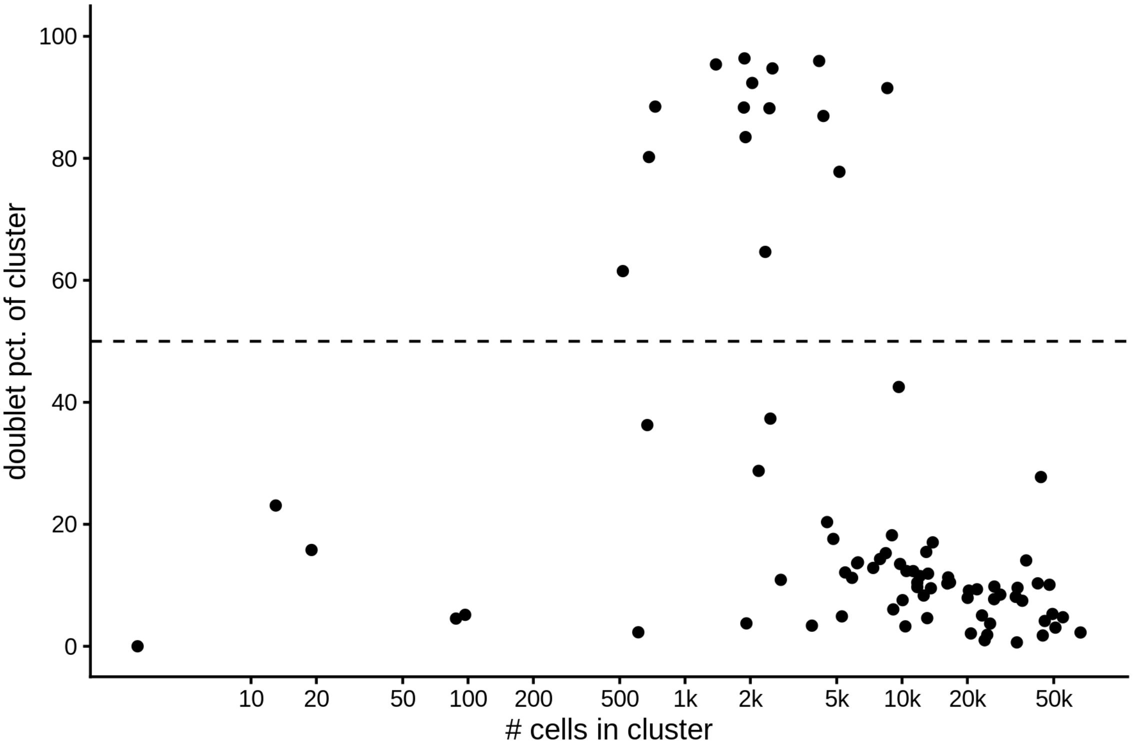

#### Clusters over UMAP

Clustering of data is performed at different resolution values. For each value the clusters are displayed over a UMAP together with a plot showing the density of cells.

After removing doublets, there were **1100463 QC-ed cells used for integration**.

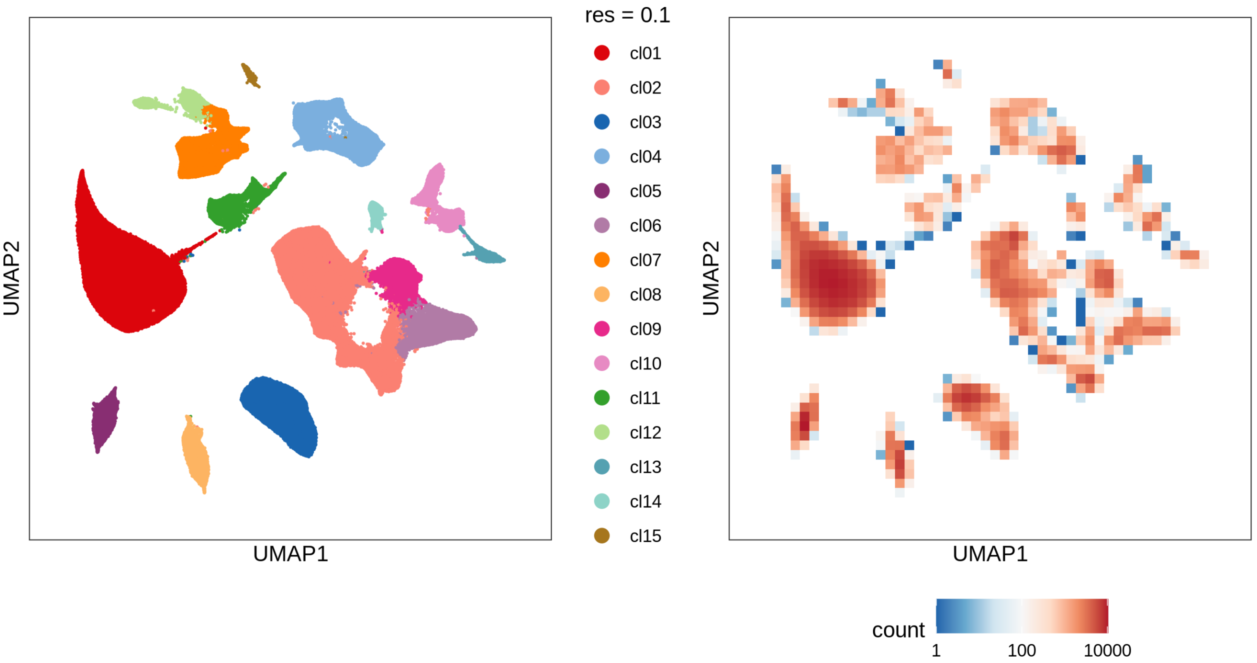

#### Evaluating cluster distribution across samples

This plot visualizes the relationship between the entropy of clusters and the maximum proportion of cells from a single sample within each cluster (higher values suggest that a cluster predominantly contains cells from a single sample). Entropy measures how evenly distributed cells are across samples within each cluster—higher entropy indicates that cells from different samples are more evenly distributed, while lower entropy suggests that a cluster is dominated by cells from a small number of samples.

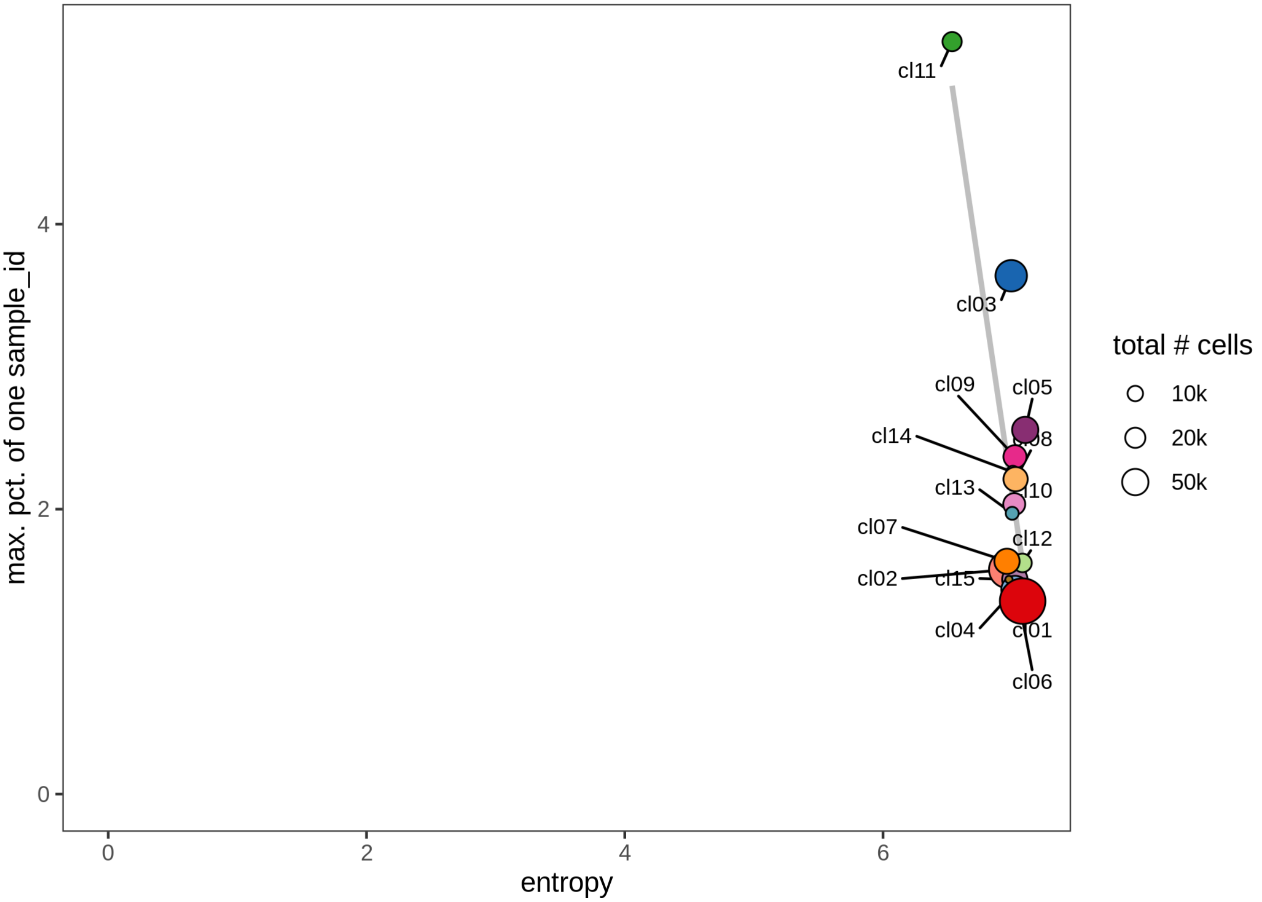

#### Check QC metrics of clusters

Distributions of QC metrics (library size, number of features, mitochondrial proportion, and spliced proportion) are shown for each cluster across different resolution values.

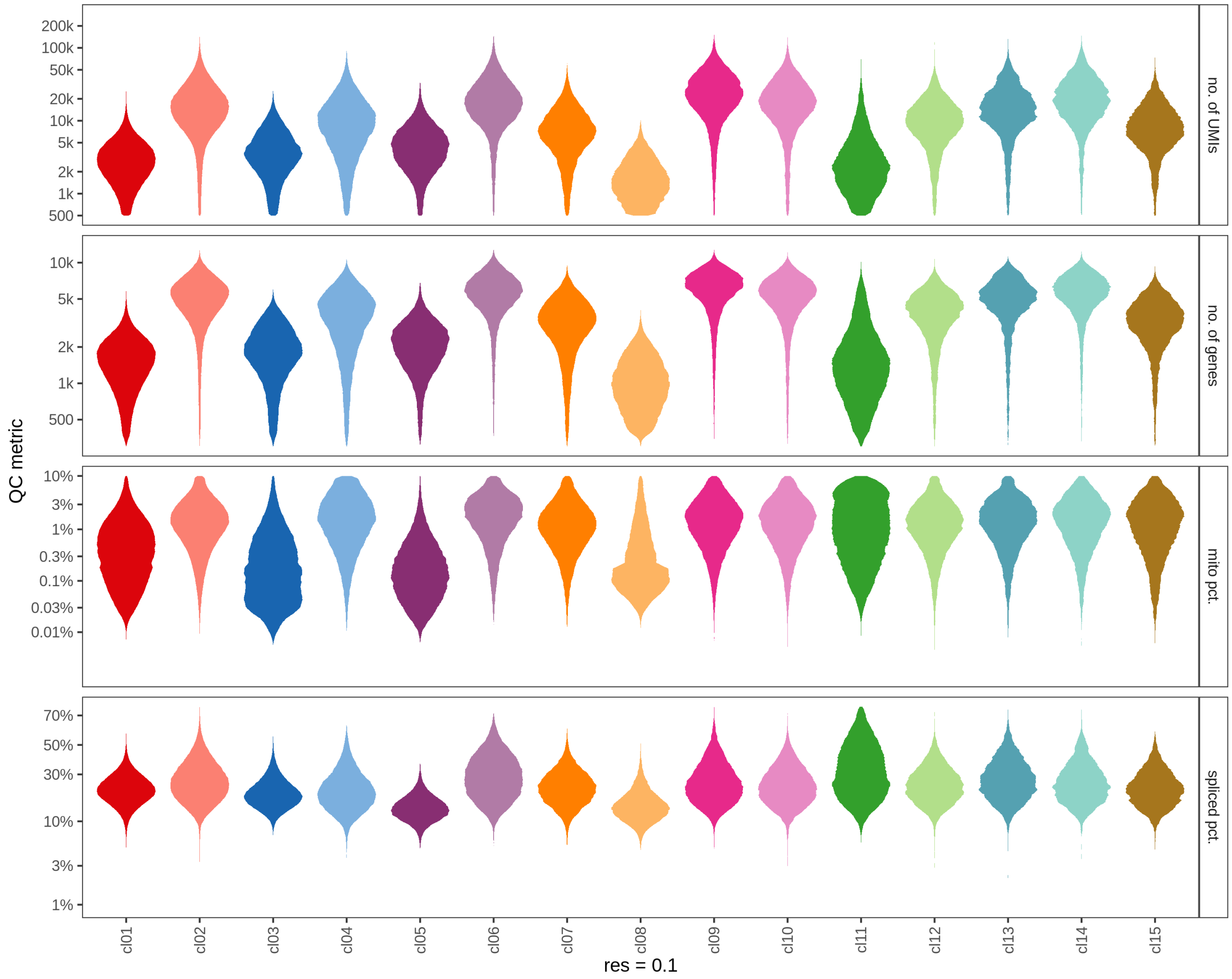

QC metric
200k
100k
50k
20k
10k
5k
2k
1k
500
no. of UMIs
10k
5k
2k
1k
500
no. of genes
10%
3%
1%
0.3%
0.1%
0.03%
0.01%
mito pct.
70%
50%
30%
10%
3%
1%
spliced pct.
cl01
cl02
cl03
cl04
cl05
cl06
cl07
cl08
cl09
cl10
cl11
cl12
cl13
cl14
cl15
res = 0.1

### Marker genes

#### Clusters over UMAP

Clustering of data is performed at resolution 0.2. Cluster membership of cells is displayed on a UMAP.

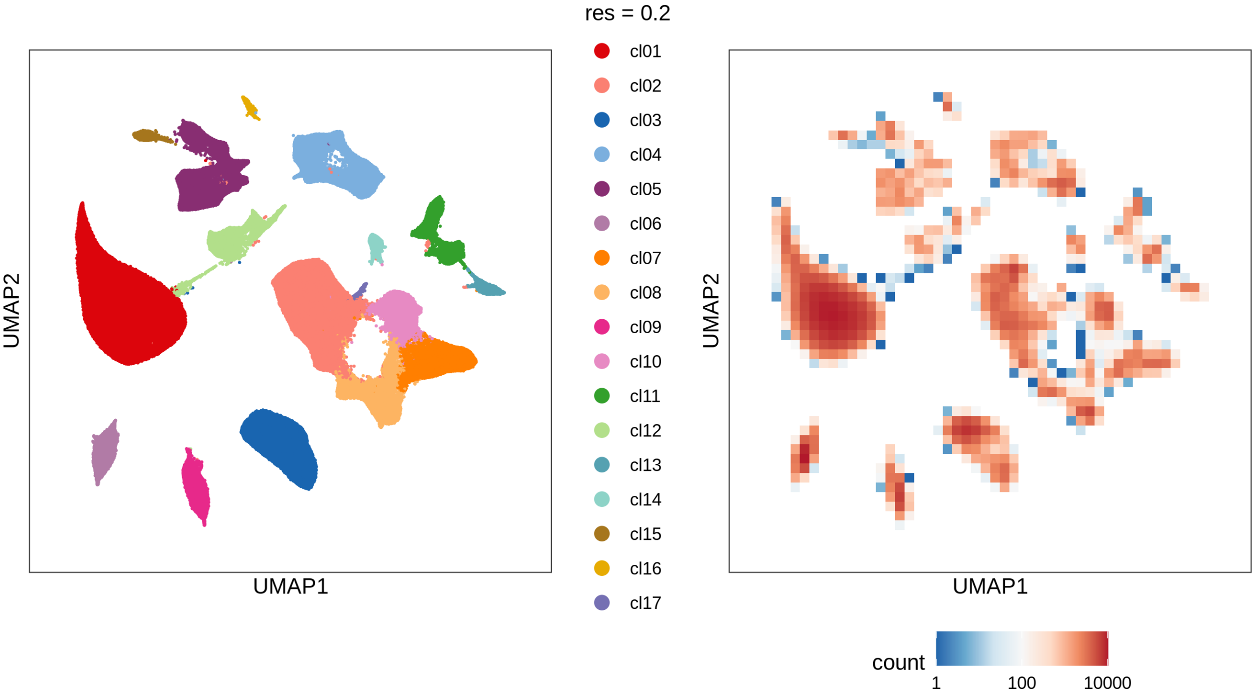

#### Cluster splits by metadata variables

For each cluster the proportion of cells coming from samples associated with specific values of diagnosis, diagnosis_subtype and original_tissue_annotation is shown.

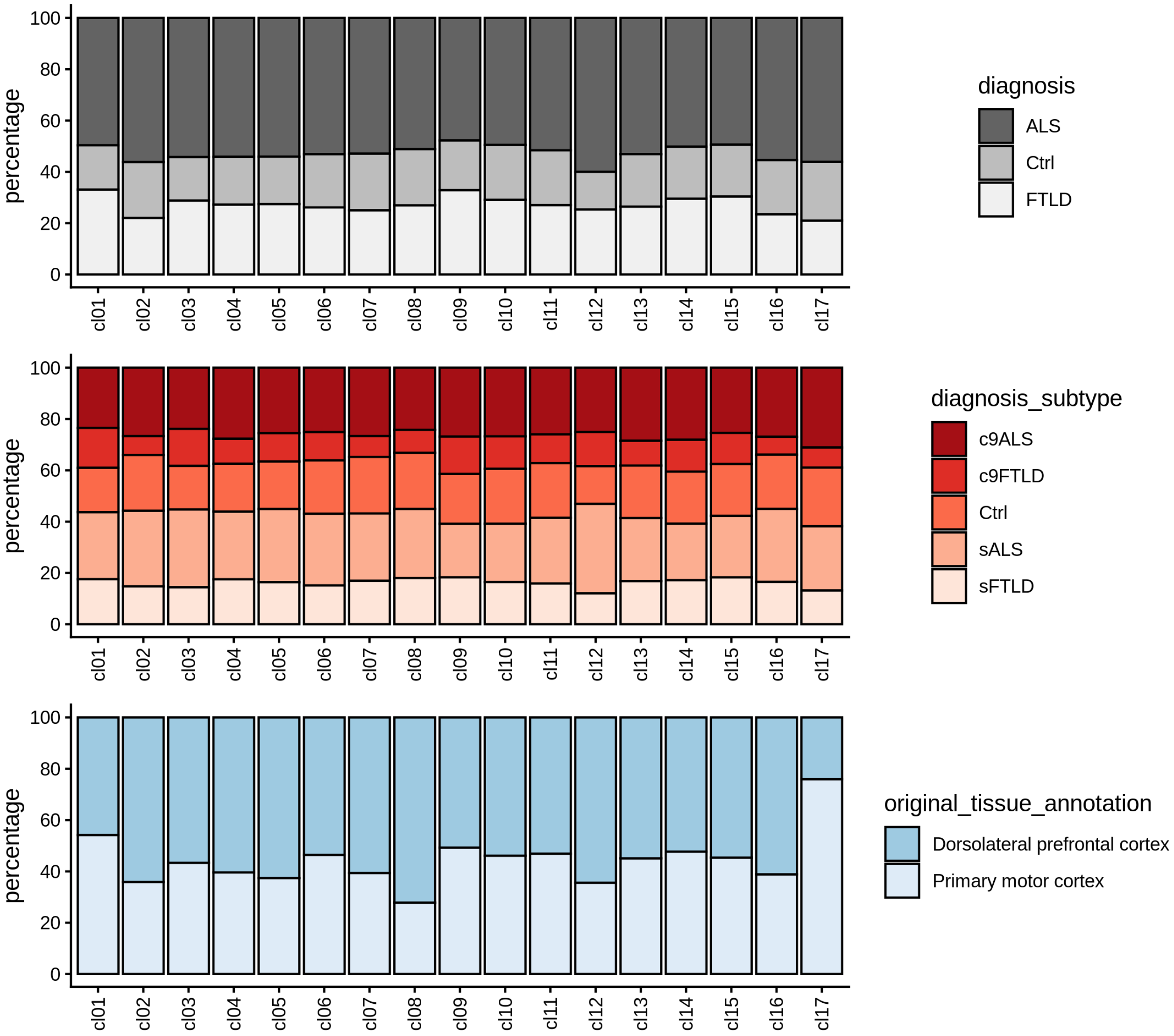

#### Metadata variables over UMAP

The plot shows a binned UMAP with facets corresponding to specific values of different metadata variables which allows the evaluation of whether cells sharing certain annotations are particularly abundant in some clusters.

```
## ### diagnosis
```

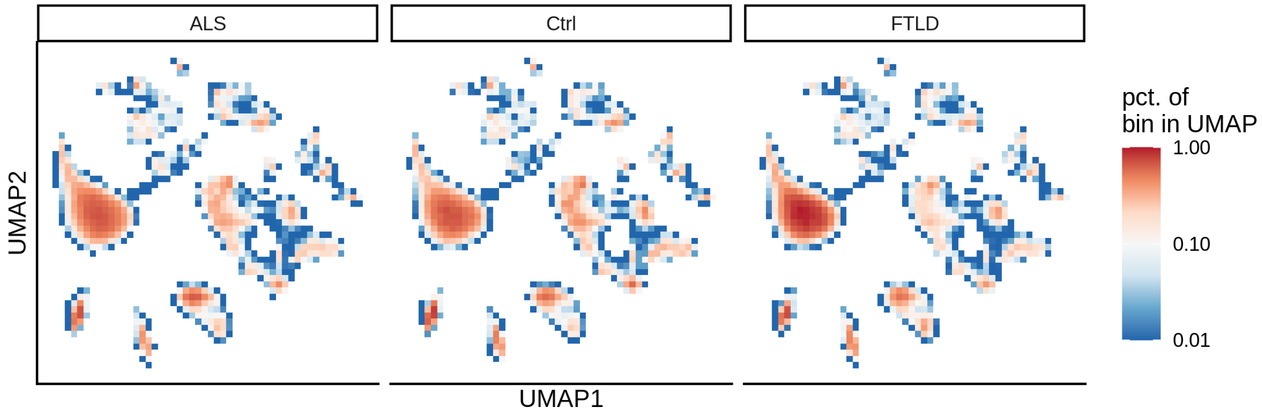

#### Marker genes

To identify marker genes for each cluster, pseudobulk counts are generated by aggregating the expression values of cells within a specific cluster for each sample. Similarly, pseudobulk counts are generated for the remaining clusters by aggregating expression values separately for each sample. The resulting pseudobulk values for the target cluster are then compared to those of the remaining clusters using `edgeR`.

##### Heatmaps of marker genes

The resulting `log2` fold change values (calculated using `edgeR`) per cluster are shown for several genesets:

- selected canonical marker genes, if specified in the config file

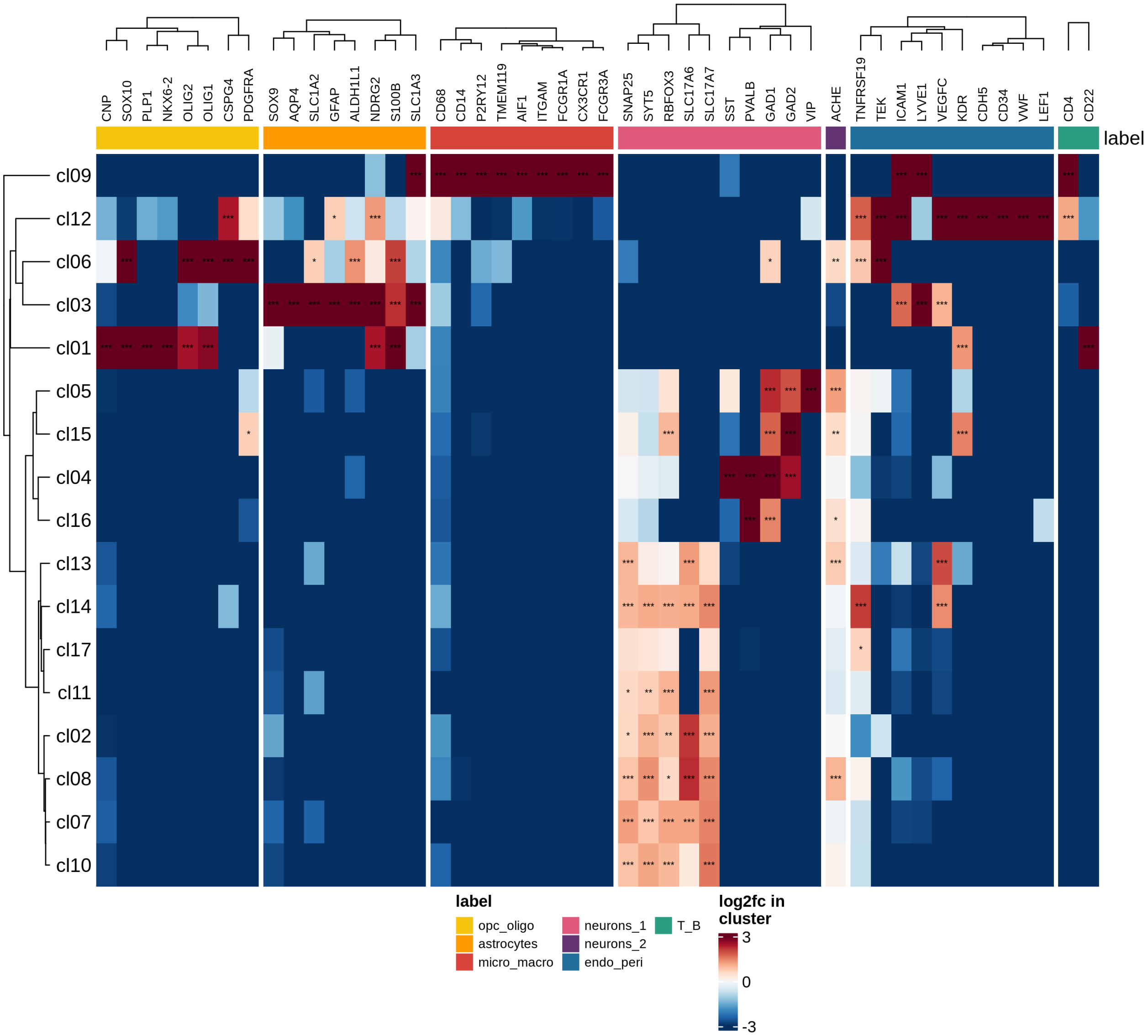

- the 50 most highly variable genes

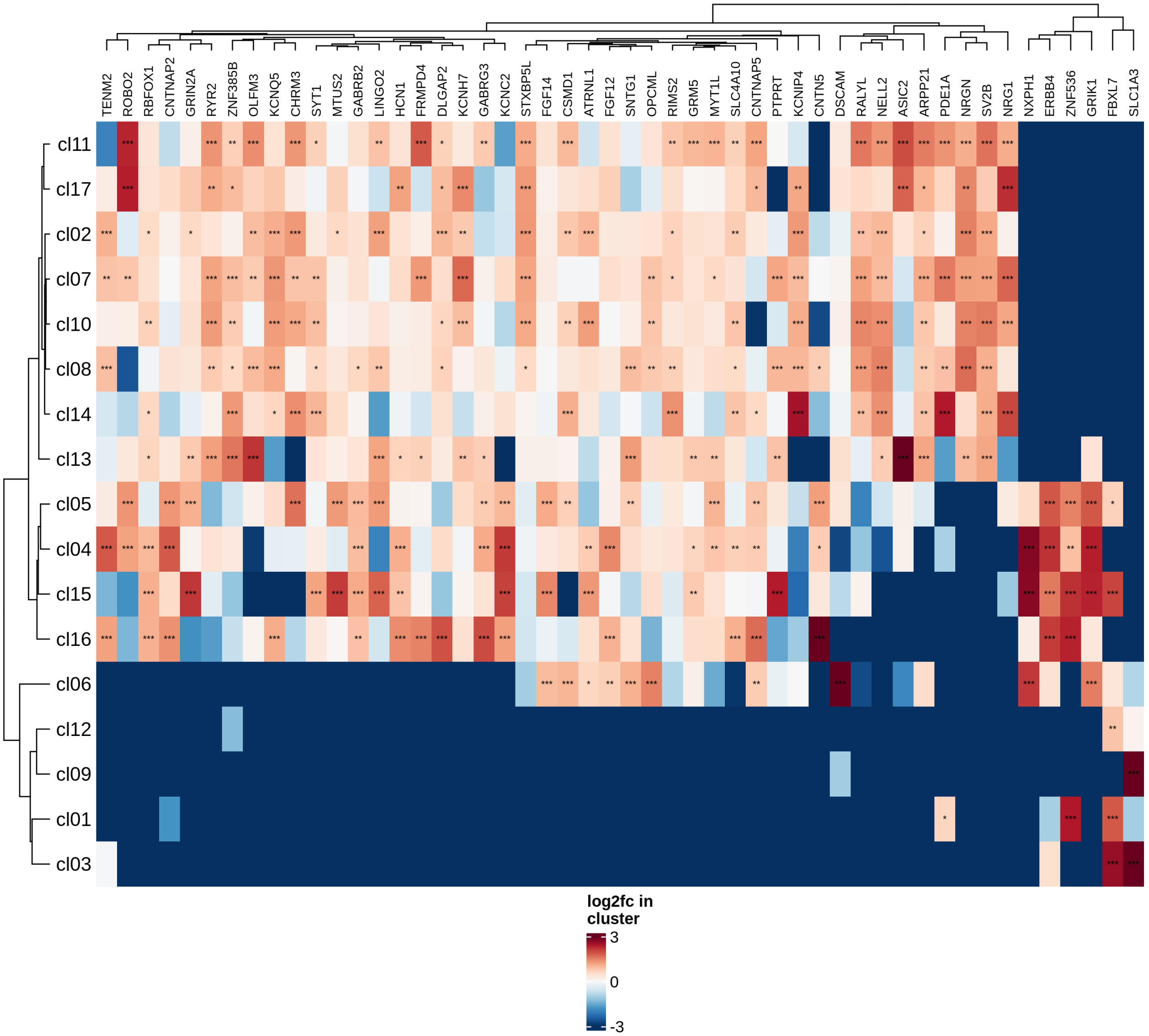

- the 50 genes most likely to represent ambient RNA contamination.

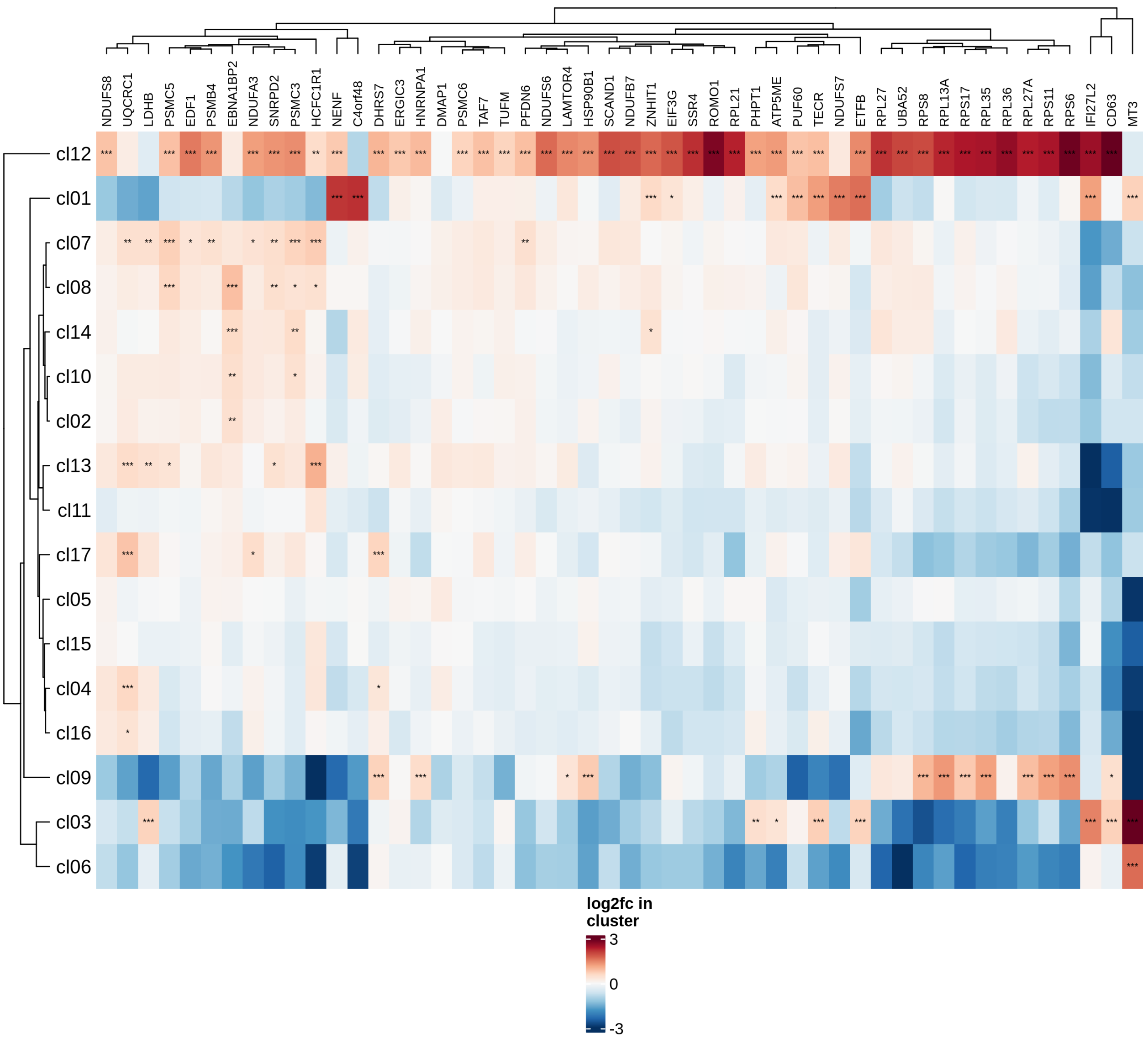

##### Highly variable genes

Normalized pseudobulk expression values for top 100 highly variable genes are shown for each cluster, in descending order of variance (variance calculated using `DESeq2::vst`).

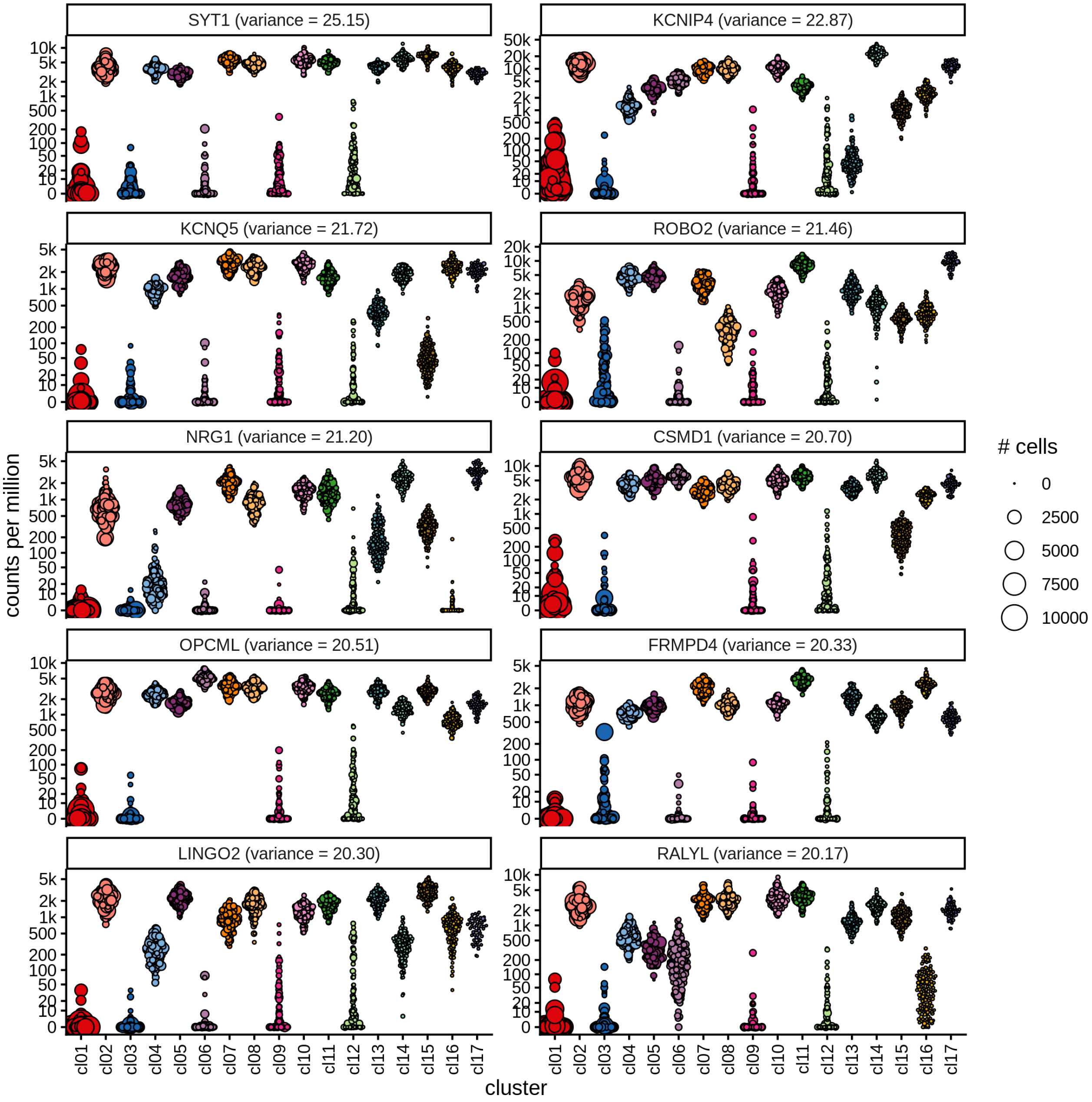

##### Top marker genes

Normalized pseudobulk expression values of top 10 marker genes with FDR < 0.05 and a minimum expression of 50 CPM are shown for each cluster.

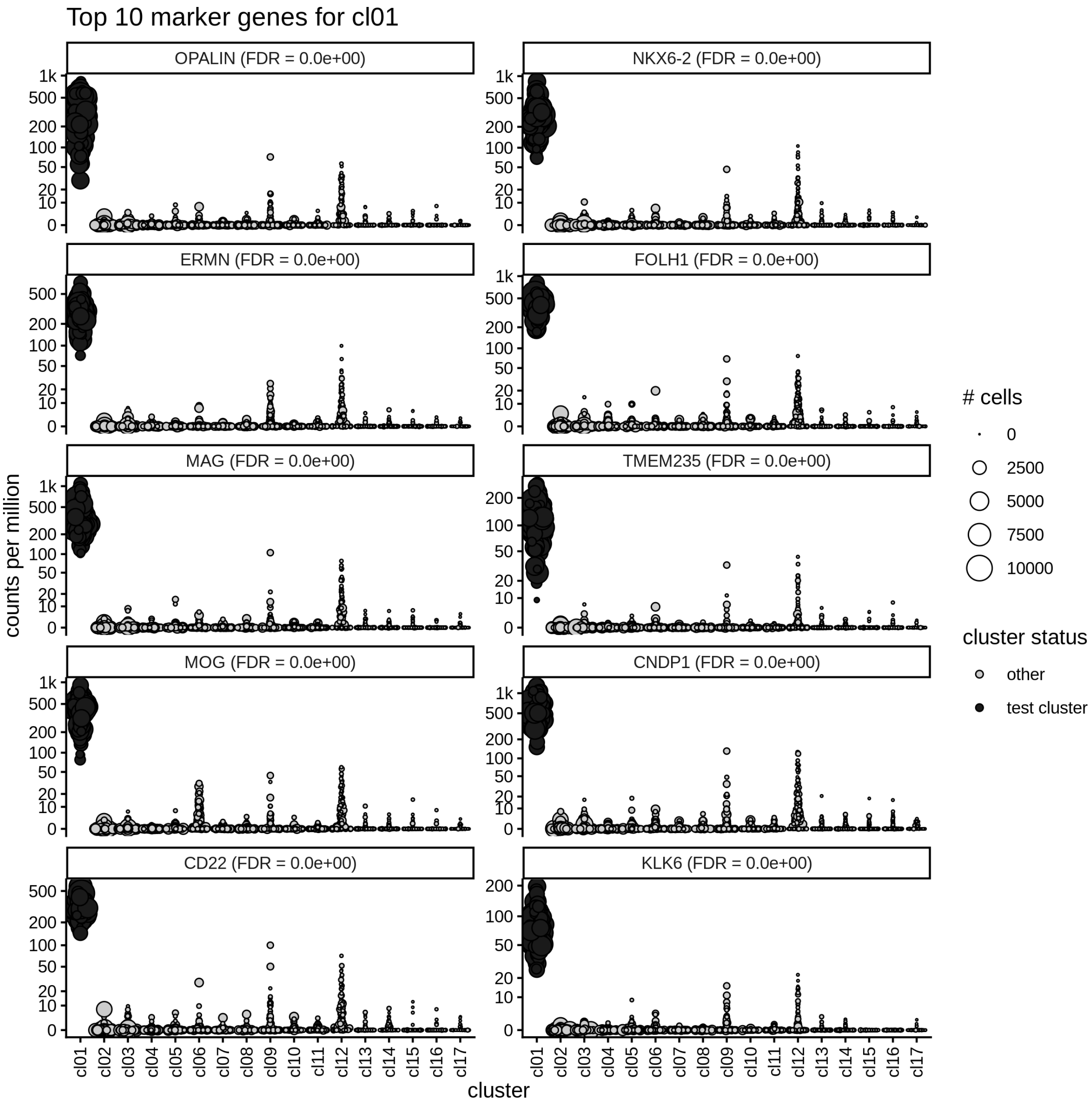

#### GSEA characterisation of clusters

Gene Set Enrichment Analysis (GSEA) was performed on marker genes for each cluster, using log fold change as the ranking variable. The top 10 pathways, grouped into five categories and selected based on a significance threshold of 0.05, are displayed for each cluster.

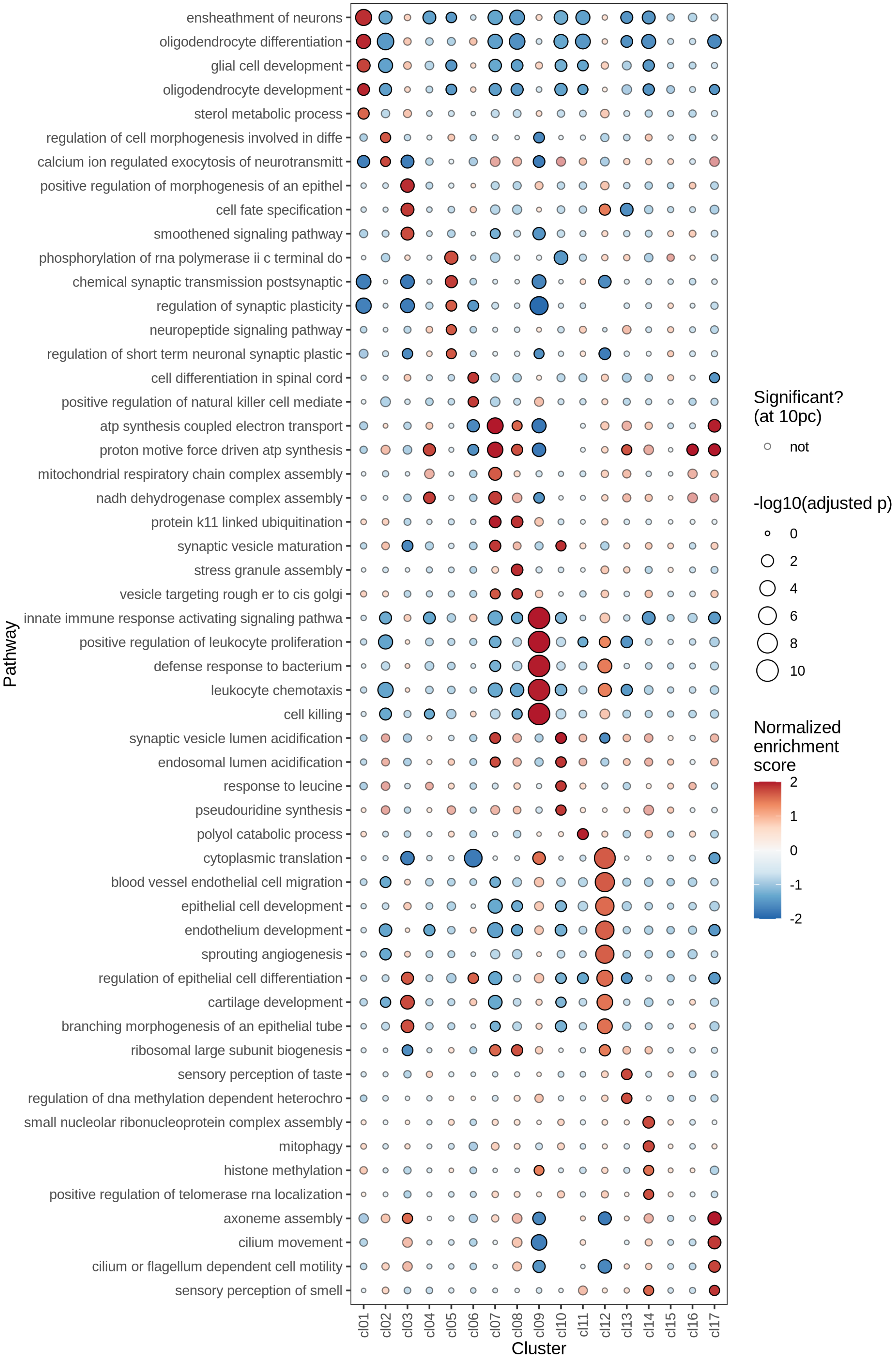
ensheathment of neurons
oligodendrocyte differentiation
glial cell development
oligodendrocyte development
sterol metabolic process
regulation of cell morphogenesis involved in diffe
calcium ion regulated exocytosis of neurotransmitt
positive regulation of morphogenesis of an epithel
cell fate specification
smoothened signaling pathway
phosphorylation of rna polymerase ii c terminal do
chemical synaptic transmission postsynaptic
regulation of synaptic plasticity
neuropeptide signaling pathway
regulation of short term neuronal synaptic plastic
cell differentiation in spinal cord
positive regulation of natural killer cell mediate
atp synthesis coupled electron transport
proton motive force driven atp synthesis
mitochondrial respiratory chain complex assembly
nadh dehydrogenase complex assembly
protein k11 linked ubiquitination
synaptic vesicle maturation
stress granule assembly
vesicle targeting rough er to cis golgi
innate immune response activating signaling pathwa
positive regulation of leukocyte proliferation
defense response to bacterium
leukocyte chemotaxis
cell killing
synaptic vesicle lumen acidification
endosomal lumen acidification
response to leucine
pseudouridine synthesis
polyol catabolic process
cytoplasmic translation
blood vessel endothelial cell migration
epithelial cell development
endothelium development
sprouting angiogenesis
regulation of epithelial cell differentiation
cartilage development
branching morphogenesis of an epithelial tube
ribosomal large subunit biogenesis
sensory perception of taste
regulation of dna methylation dependent heterochro
small nucleolar ribonucleoprotein complex assembly
mitophagy
histone methylation
positive regulation of telomerase rna localization
axoneme assembly
cilium movement
cilium or flagellum dependent cell motility
sensory perception of smell
Pathway
cl01 cl02 cl03 cl04 cl05 cl06 cl07 cl08 cl09 cl10 cl11 cl12 cl13 cl14 cl15 cl16 cl17
Cluster
Significant?
(at 10pc)
not
-log10(adjusted p)
0
2
4
6
8
10
Normalized
enrichment
score
2
1
0
-1
-2

### Cell type labelling

#### Integration clusters vs predicted cell types

Prediction of cell type labels is a two-step process: The labeller first assigns a predicted cell type to each individual cell. These cell-level predictions are then aggregated at the level of high-resolution clusters (a majority-voting approach is used, where the final, assigned label for a cluster is the most common predicted label among all the cells within it).

The heatmap helps to assess whether this aggregation step seems sensible. Each cell on the heatmap shows the proportion of cells within a cluster (on the x-axis) that are assigned a specific predicted cell type in the first step (on the y-axis).

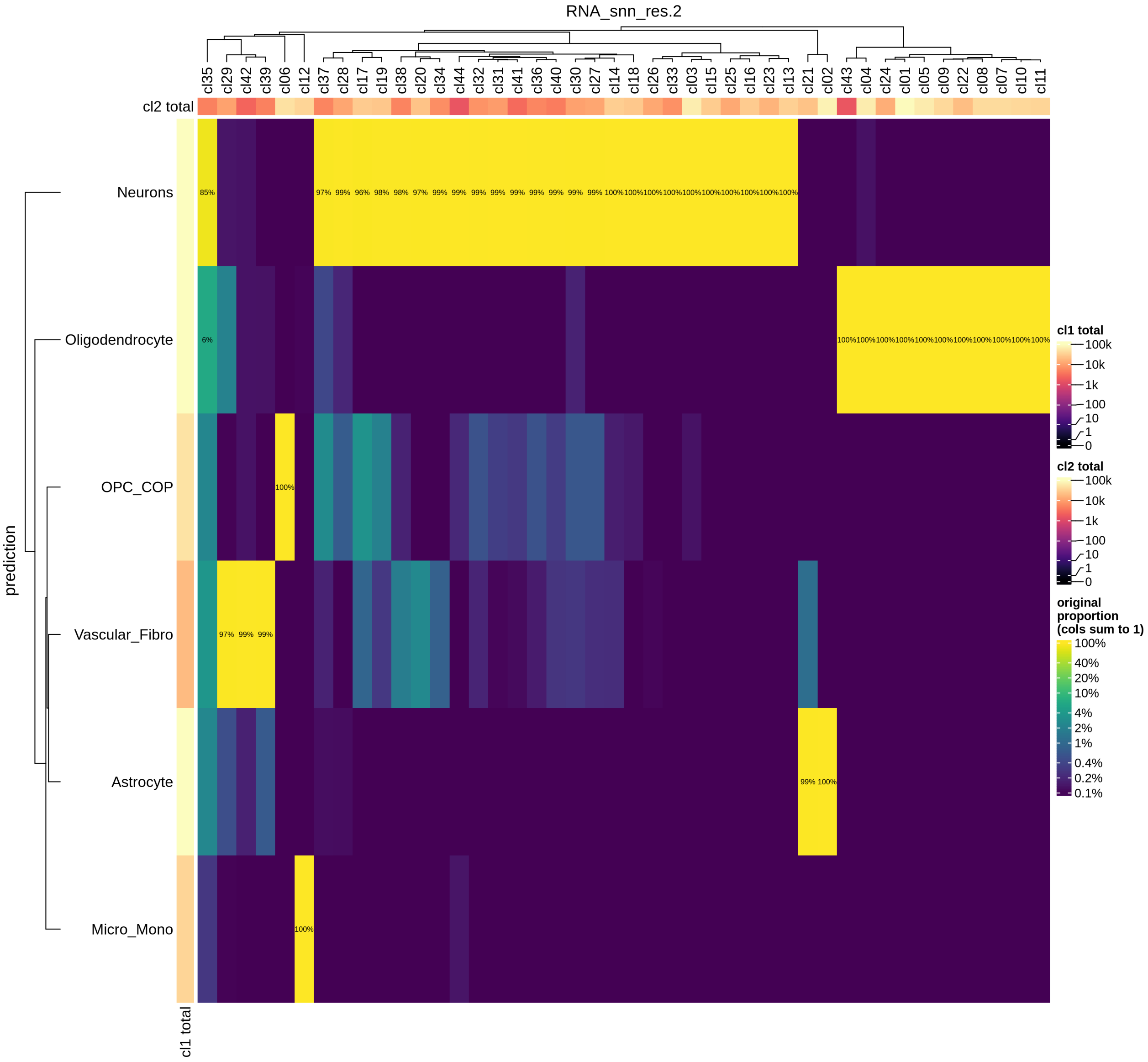

#### Predicted cell types over UMAP

Clustering of data is performed with different labellers and models. Each of these is shown over a UMAP together with a plot showing predicted cell type labels.

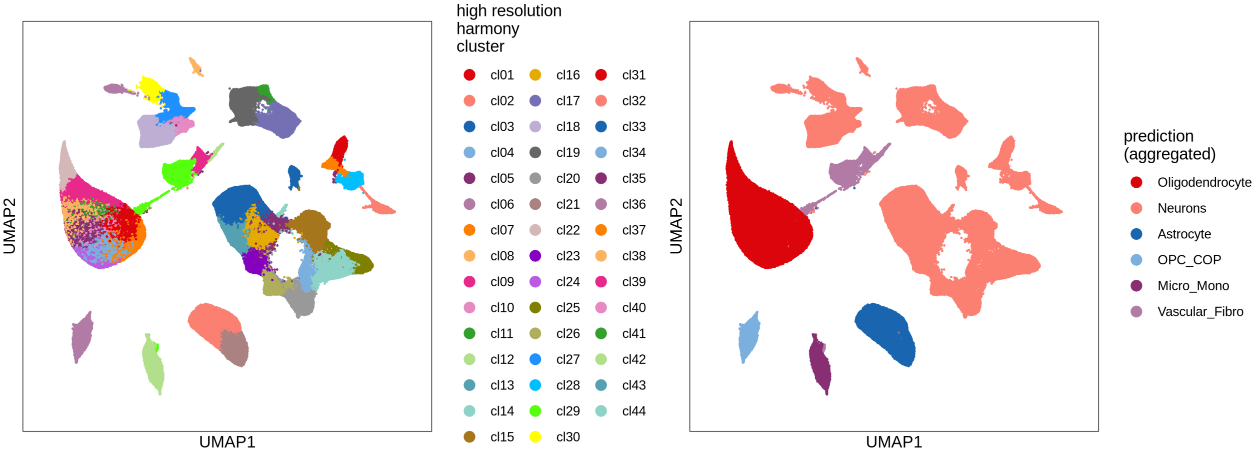
high resolution harmony cluster
cl01
cl02
cl03
cl04
cl05
cl06
cl07
cl08
cl09
cl10
cl11
cl12
cl13
cl14
cl15
cl16
cl17
cl18
cl19
cl20
cl21
cl22
cl23
cl24
cl25
cl26
cl27
cl28
cl29
cl30
cl31
cl32
cl33
cl34
cl35
cl36
cl37
cl38
cl39
cl40
cl41
cl42
cl43
cl44
UMAP1
UMAP2
prediction (aggregated)
Oligodendrocyte
Neurons
Astrocyte
OPC_COP
Micro_Mono
Vascular_Fibro
UMAP1
UMAP2

### Zoom in and recluster: myeloid

#### HVG selection diagnostics

The selection of highly variable genes is a crucial step in the integration process and in identifying clusters in single-cell experiments. However, contamination from ambient RNA can act as a significant confounding factor. This section of the report provides plots to evaluate the influence of ambient RNA on the selection of highly variable genes and its potential impact on downstream analyses.

##### Are highly variable genes “ambient” ?

The plot shows the extent to which highly variable genes identified using the `Seurat` VST method may be influenced by ambient RNA. The x-axis represents the `log2fc` values derived from the ambient gene estimation step in `scprocess` (see `scprocess` documentation for more details), while the y-axis shows the trend-normalized variance calculated using the `Seurat` VST method. Each point represents a gene, annotated based on whether it is among the top HVGs and whether it is identified as ambient by the ambient gene detection step. Labelled genes are top 10 most variable.

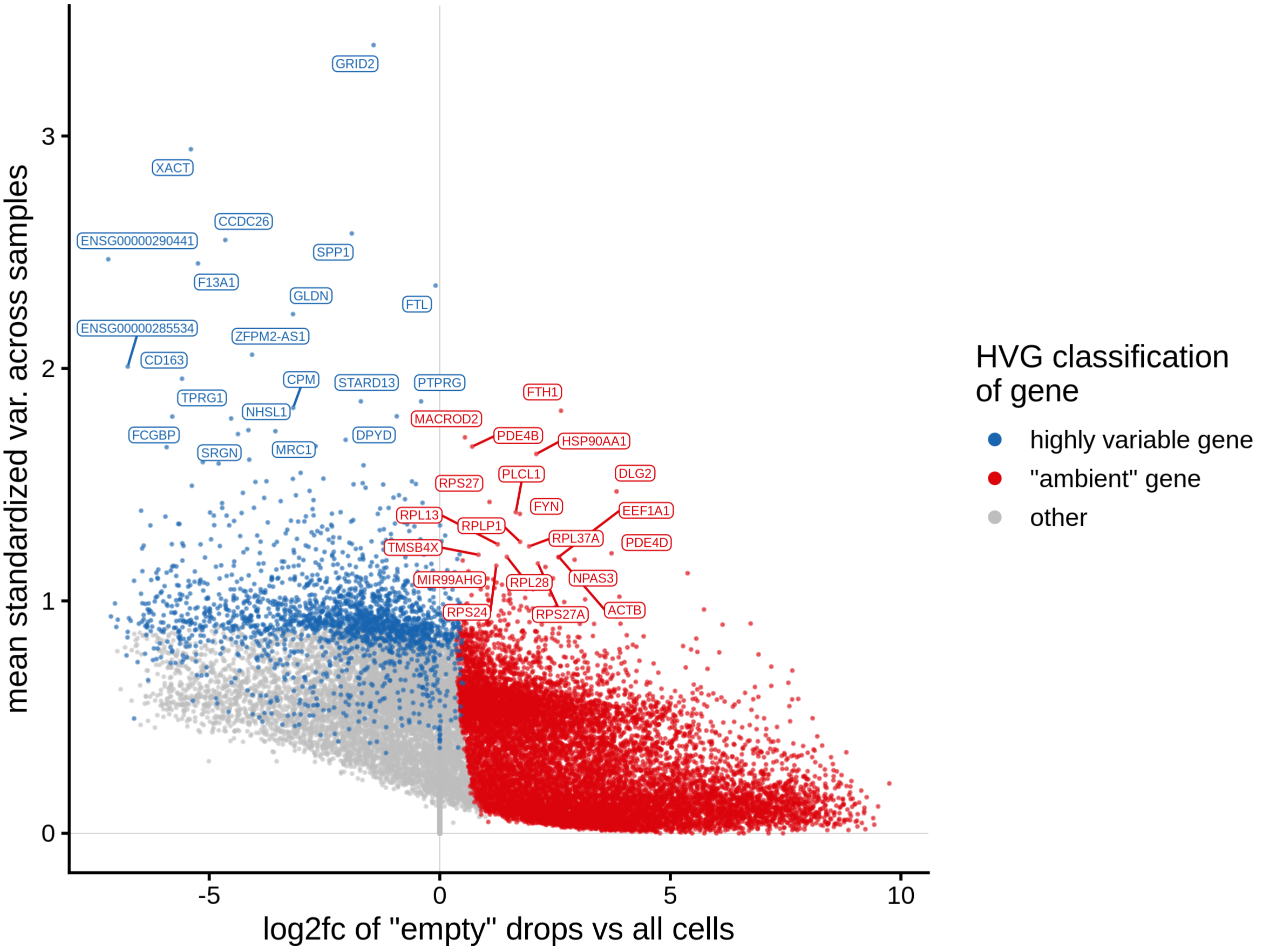

##### Which genes are “ambient”?

The plot shows the results of the `scprocess` ambient gene detection procedure. The y-axis is the `-log10` nominal p-value, with a dotted line indicating the threshold where the adjusted p-value is sufficiently small ($< 0.01$). Multiple plots are shown, each corresponding to a different minimum expression level filter applied to the ambient profiles.

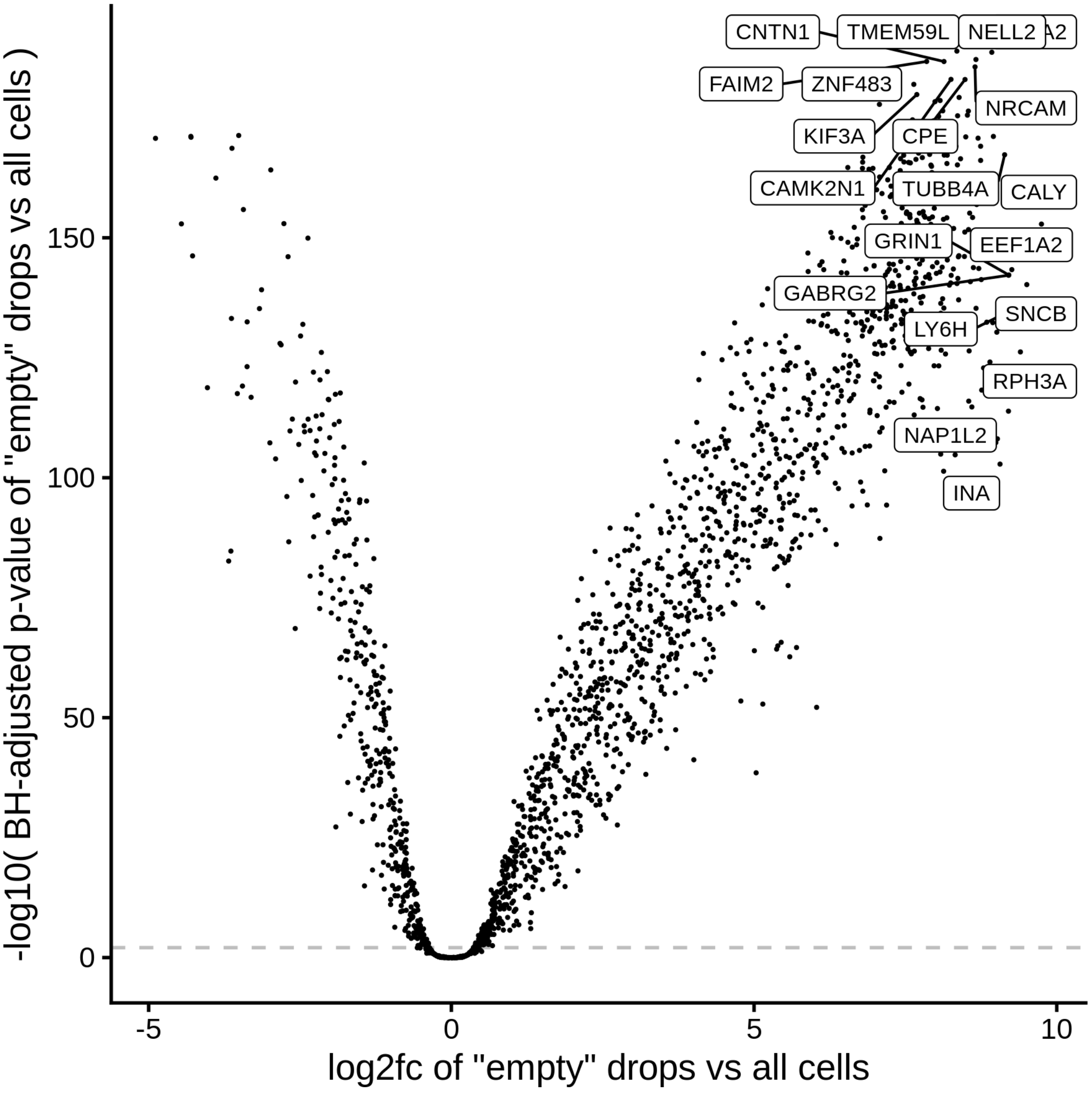

**Which genes are variable across the ambient profiles?**

Examining how ambient genes vary across different samples can provide valuable insights. Such variation may highlight cases where certain samples require distinct treatment, for example, if case and control samples consistently exhibit different ambient profiles.

The heatmaps display the pseudobulk expression of various genes across the ambient profiles of each sample in the dataset. The genes shown are the top 40 genes with highest variance across ambient profiles;

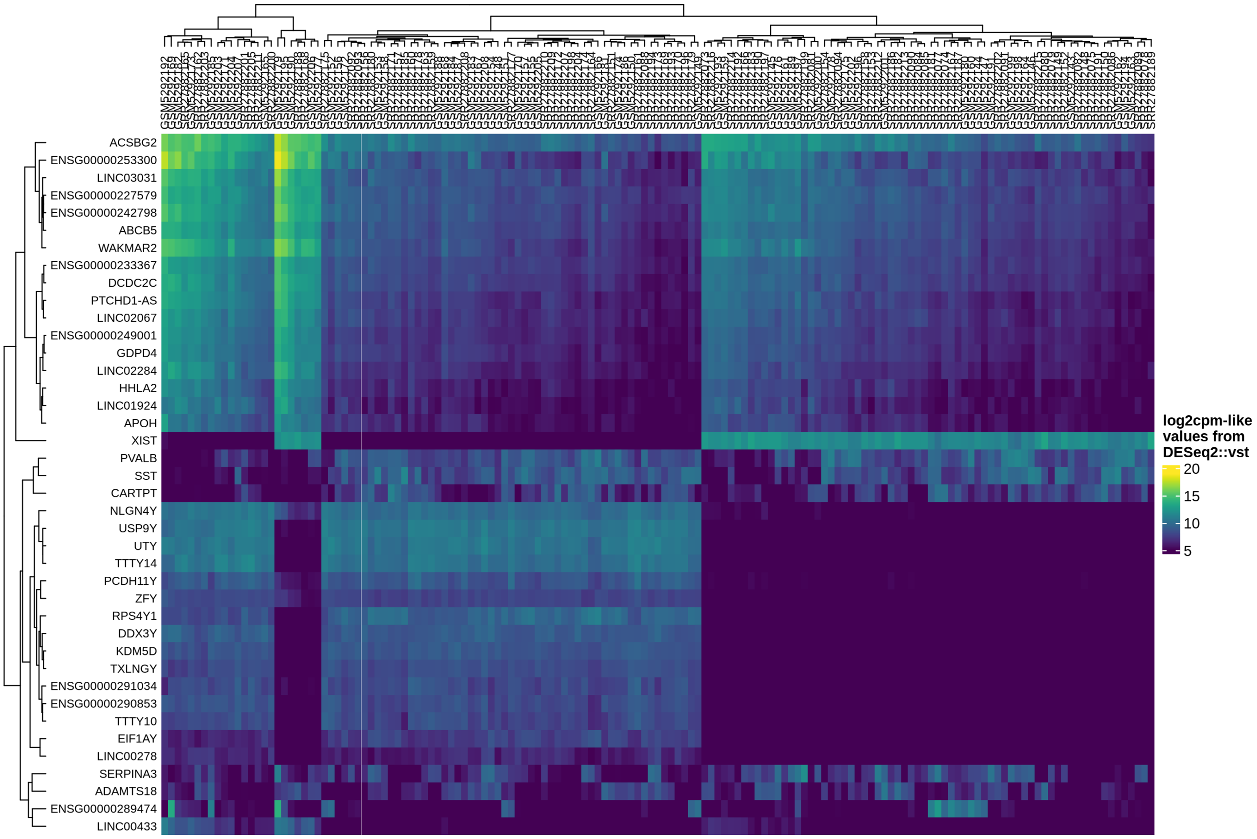

### Clustering diagnostics

#### Clusters over UMAP

Clustering of data is performed at different resolution values. For each value the clusters are displayed over a UMAP together with a plot showing the density of cells.

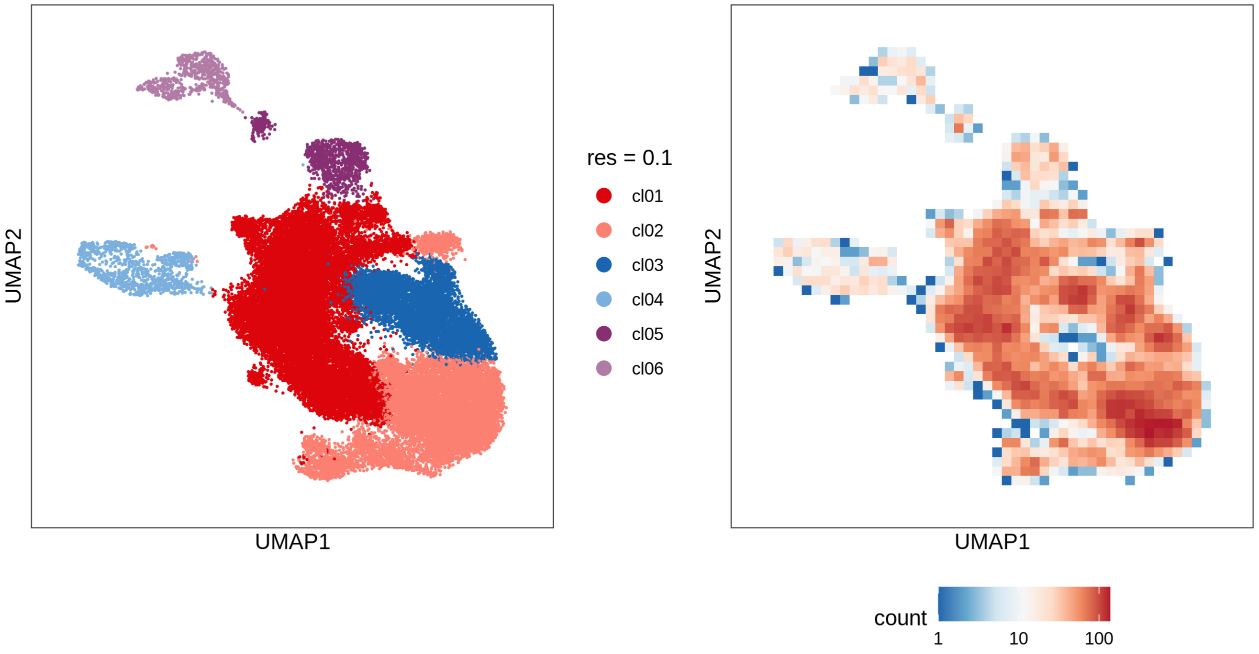

**Evaluating cluster distribution across samples**

This plot visualizes the relationship between the entropy of clusters and the maximum proportion of cells from a single sample within each cluster (higher values suggest that a cluster predominantly contains cells from a single sample). Entropy measures how evenly distributed cells are across samples within each cluster—higher entropy indicates that cells from different samples are more evenly distributed, while lower entropy suggests that a cluster is dominated by cells from a small number of samples.

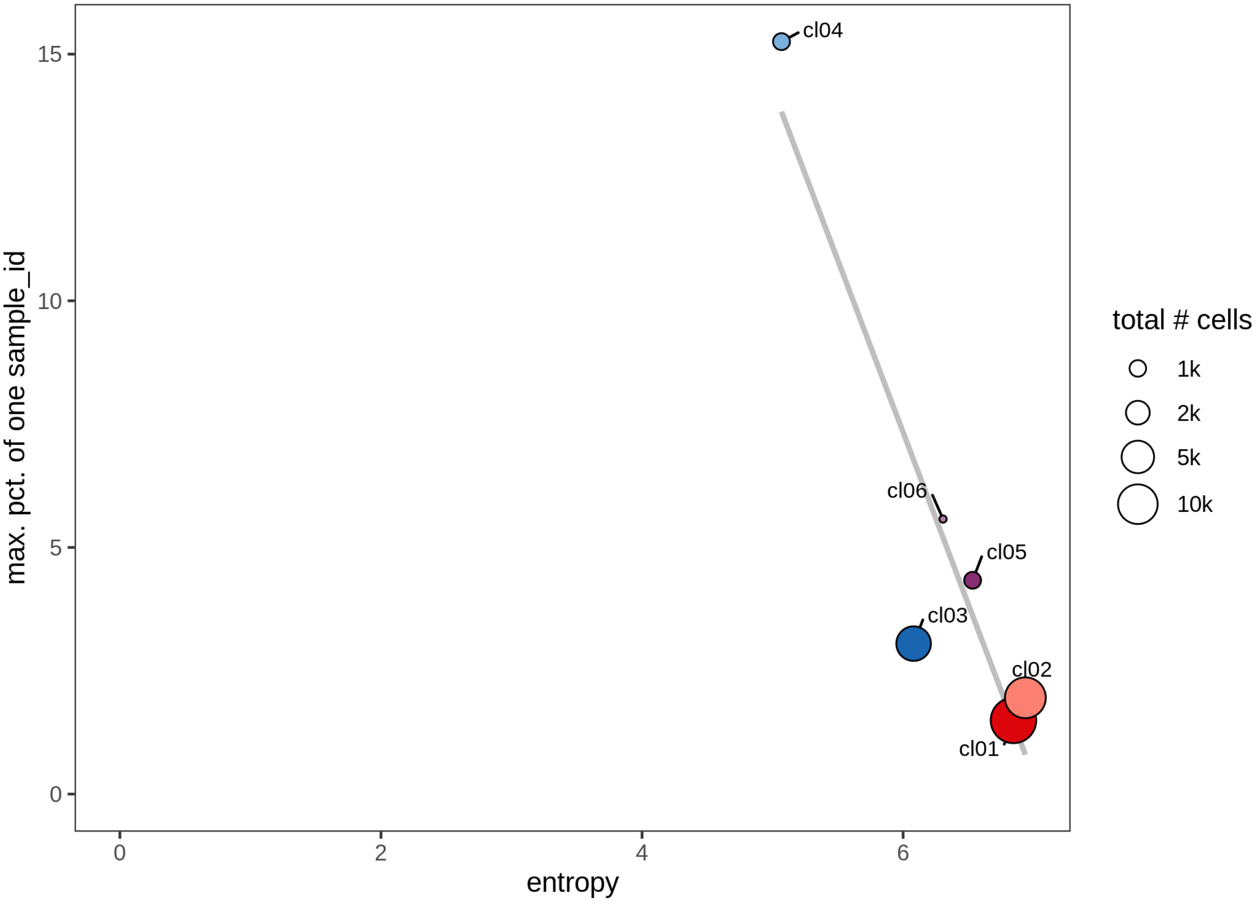

**QC metrics of clusters**

Distributions of QC metrics (library size, number of features, mitochondrial proportion, and spliced proportion) are shown for each cluster across different resolution values.

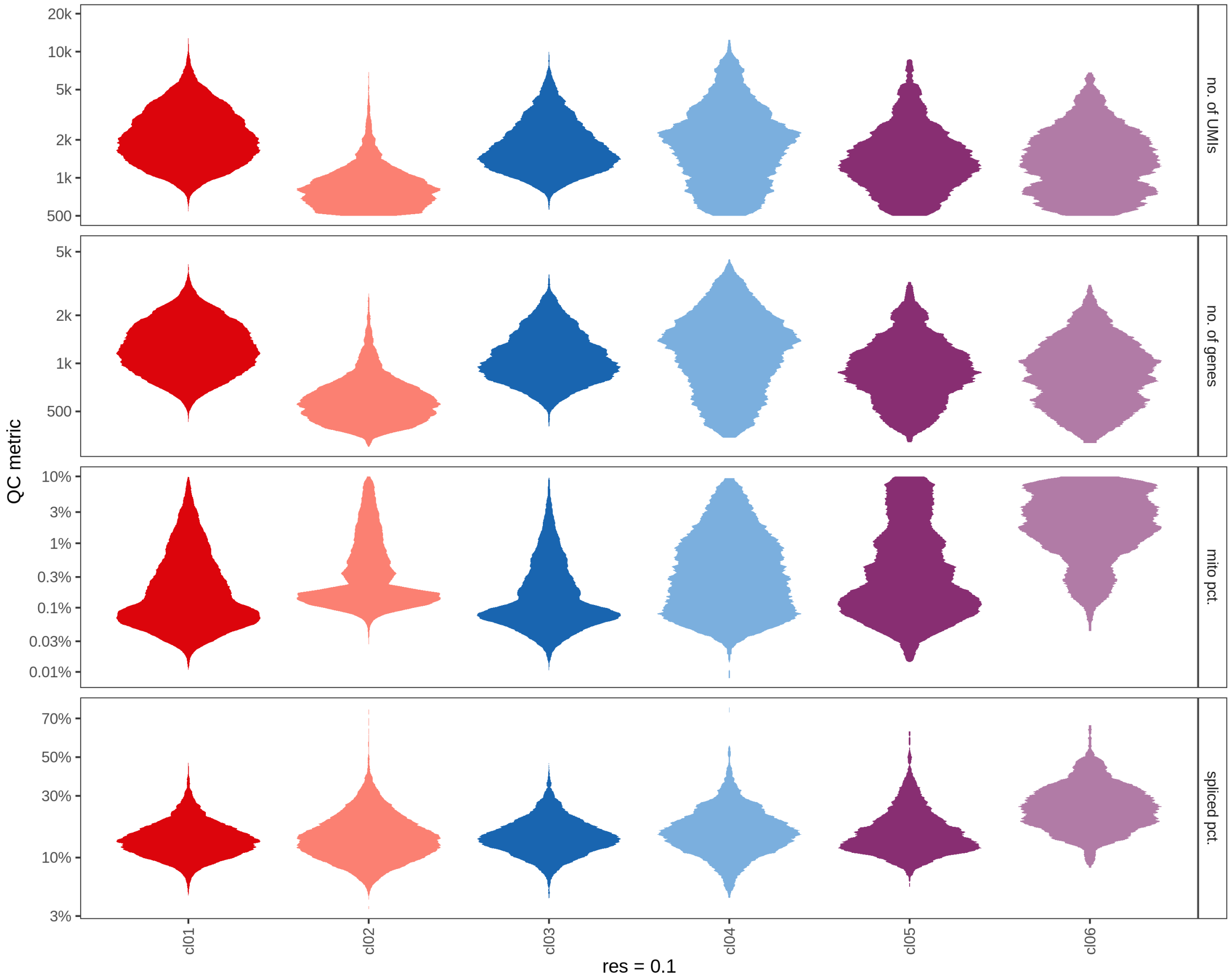

##### Cluster splits by metadata variables

For each cluster (resolution: 0.5) the proportion of cells coming from samples associated with specific values of diagnosis, diagnosis_subtype and original_tissue_annotation is shown.

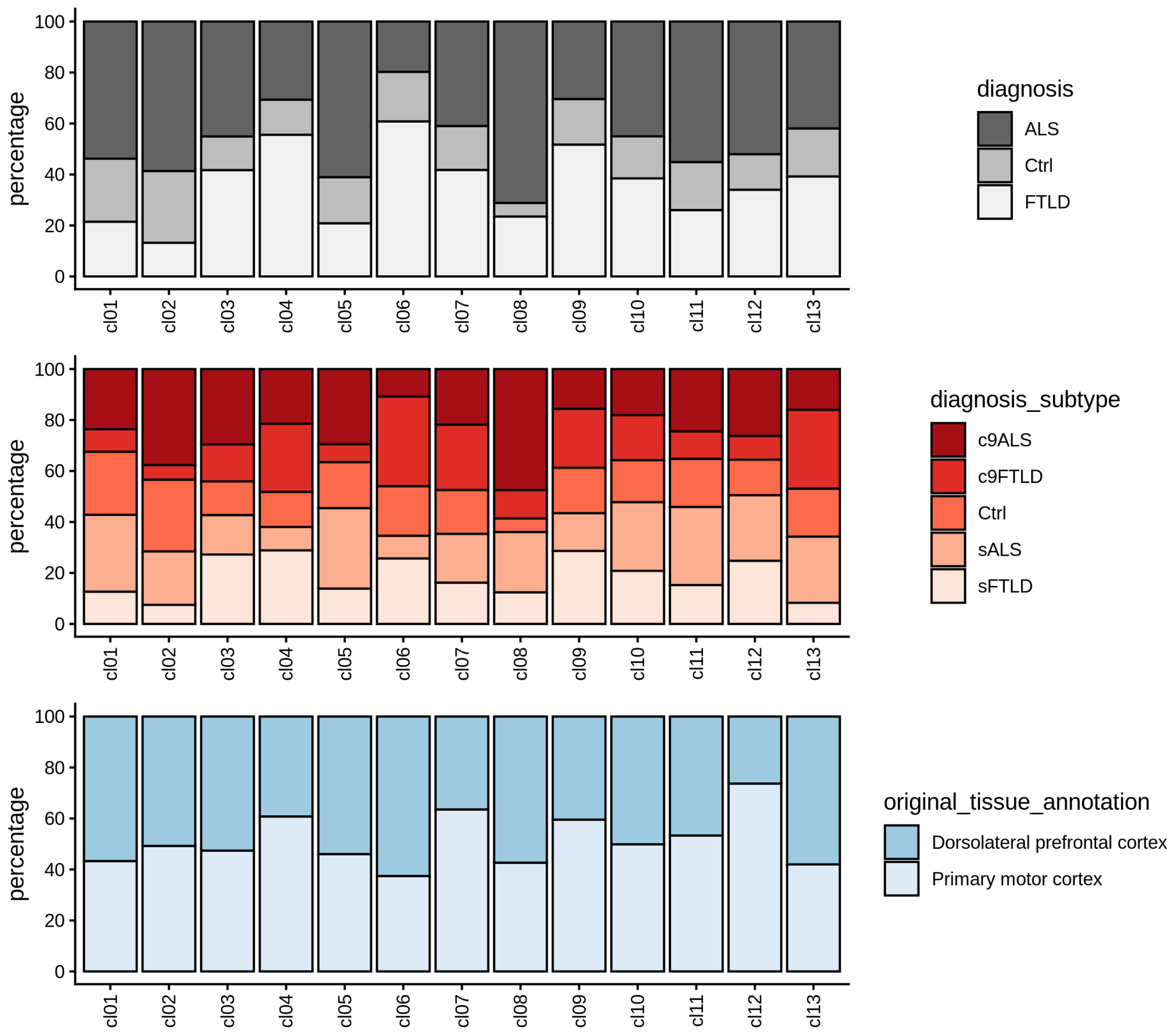

##### Metadata variables over UMAP

The plot shows a binned UMAP with facets corresponding to specific values of different metadata variables which allows the evaluation of whether cells sharing certain annotations are particularly abundant in some clusters.

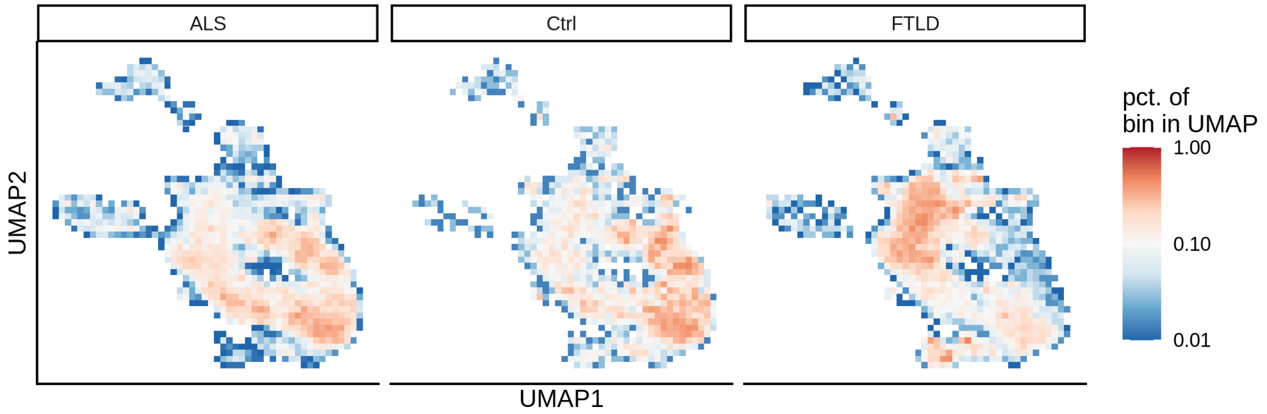

#### Marker genes

To identify marker genes for each cluster (resolution: 0.5), pseudobulk counts are generated by aggregating the expression values of cells within a specific cluster for each sample. Similarly, pseudobulk counts are generated for the remaining clusters by aggregating expression values separately for each sample. The resulting pseudobulk values for the target cluster are then compared to those of the remaining clusters using `edgeR`.

##### Heatmaps of marker genes

The resulting `log2` fold change values (calculated using `edgeR`) per cluster are shown for several the 50 most highly variable genes

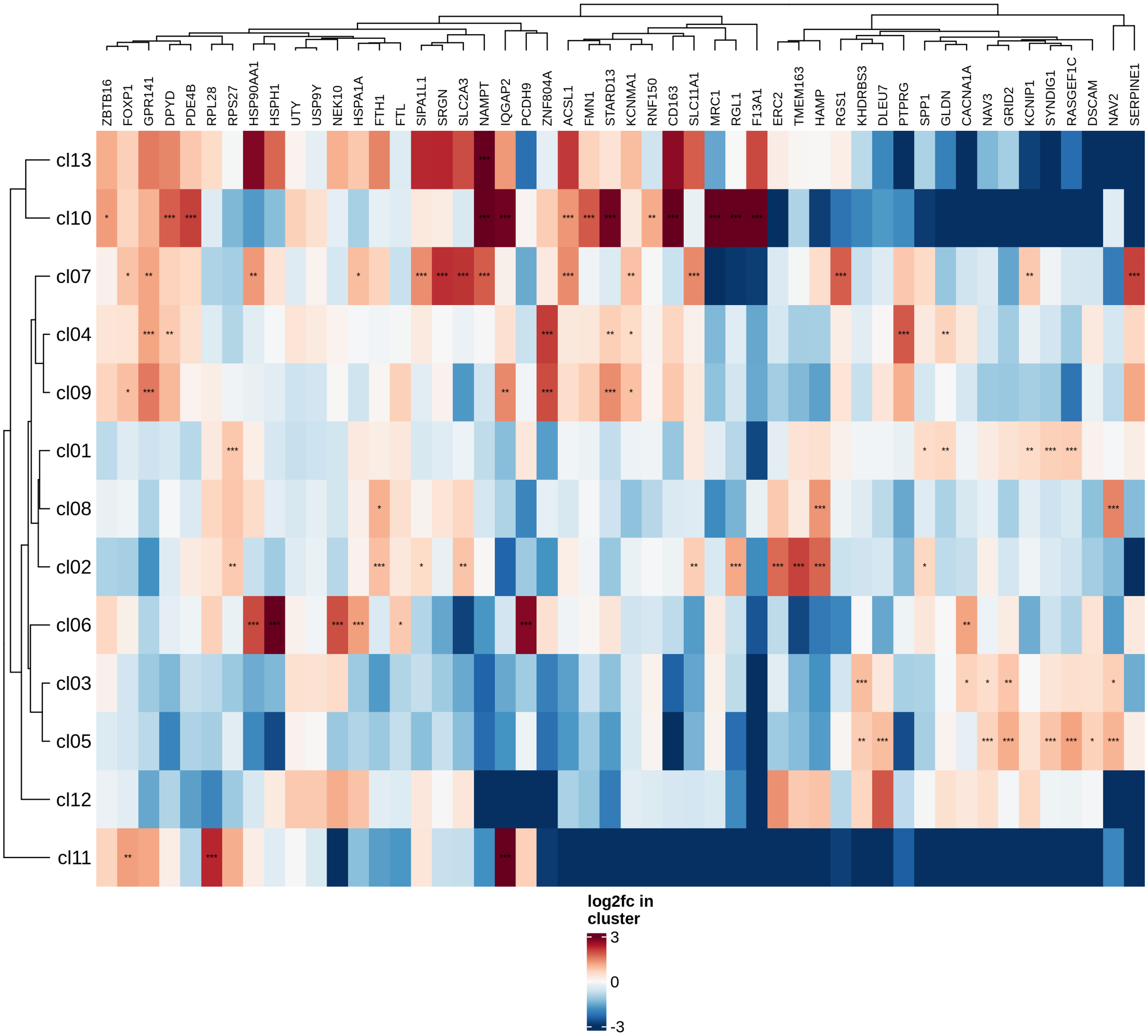

##### Highly variable genes

Normalized pseudobulk expression values for top 100 highly variable genes are shown for each cluster, in descending order of variance (variance calculated using `DESeq2::vst`).

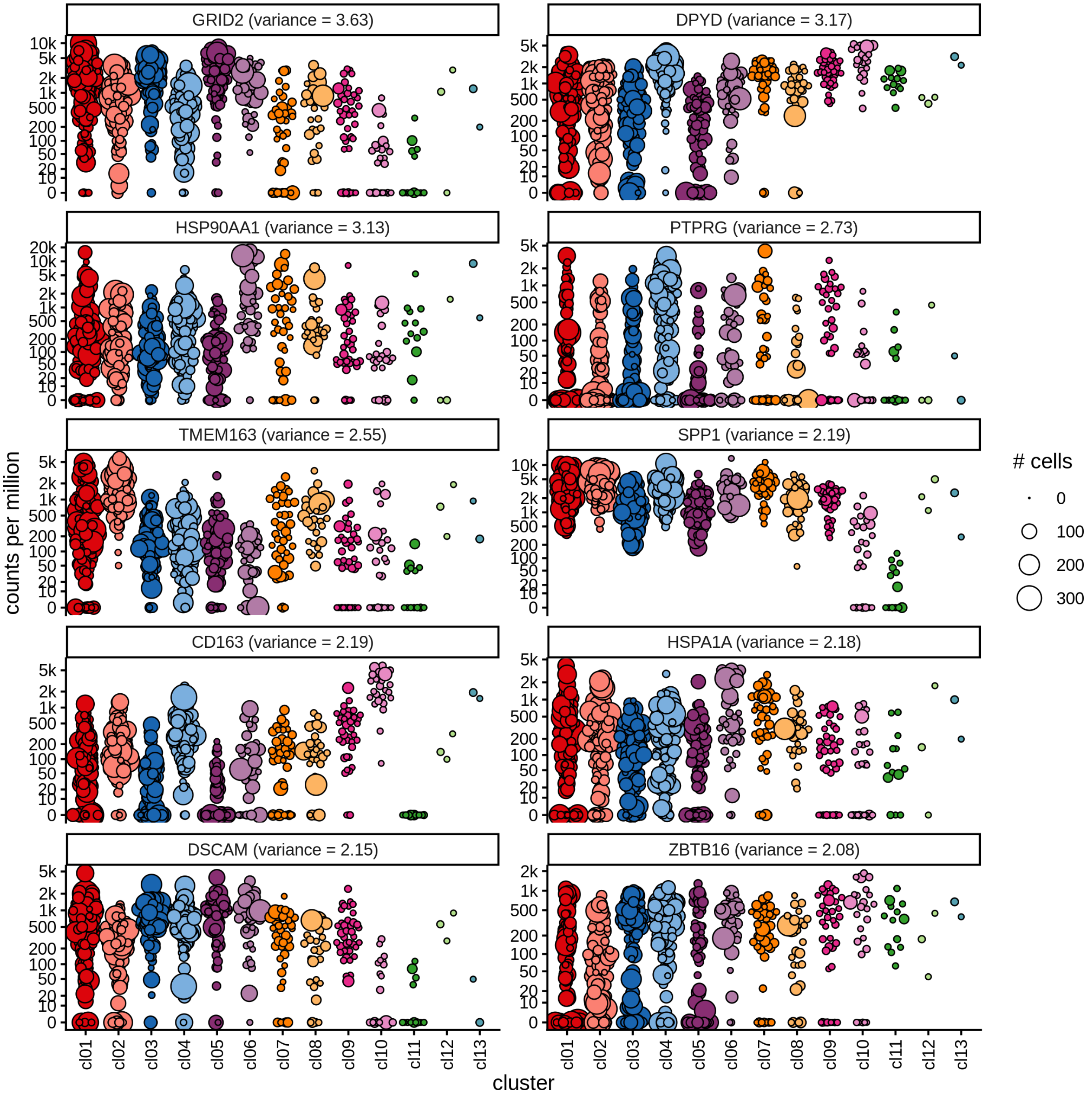

#### Top marker genes

Normalized pseudobulk expression values of top 10 marker genes with FDR < 0.05 and a minimum expression of 50 CPM are shown for each cluster.

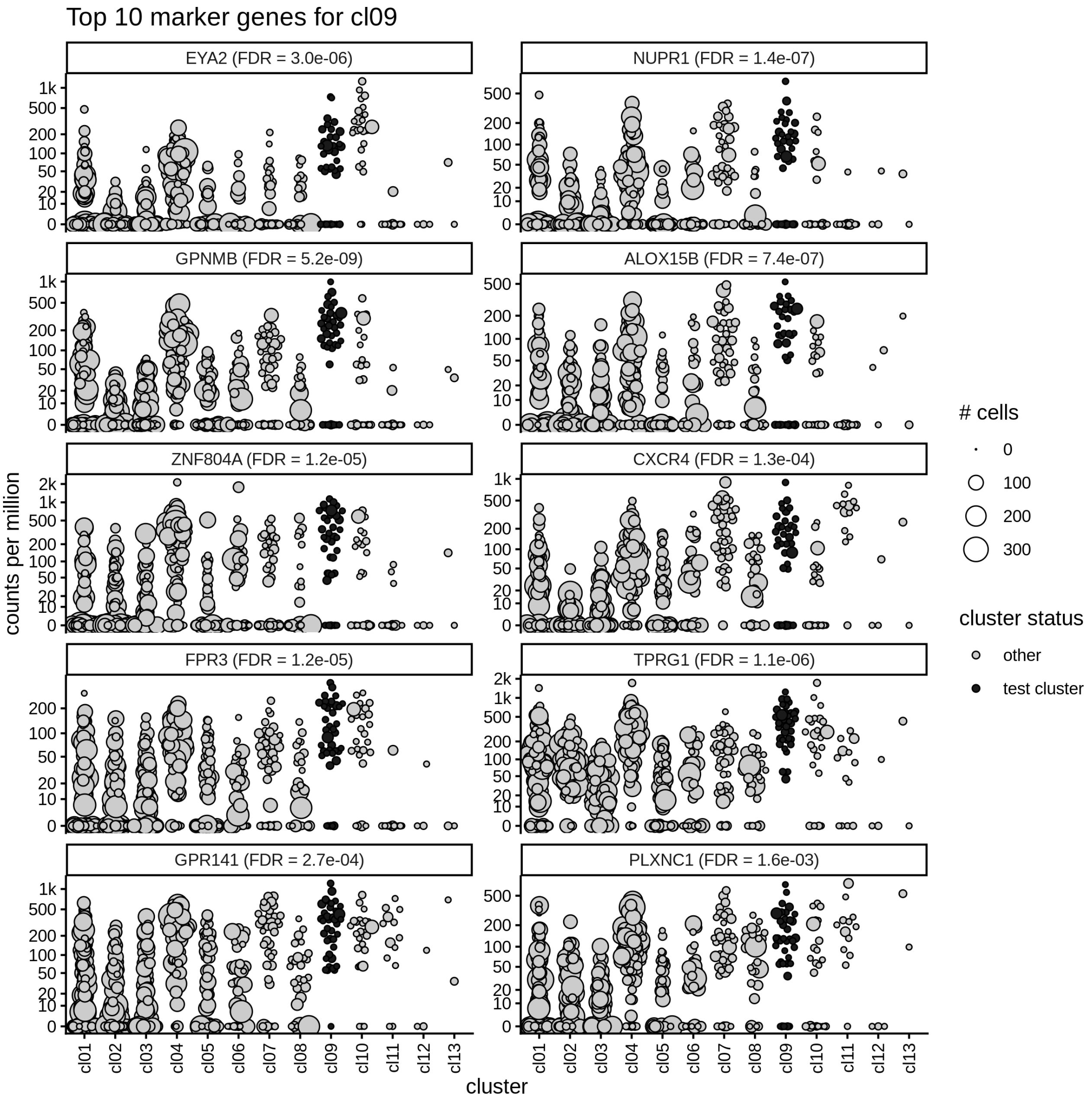
Top 10 marker genes for cl09
EYA2 (FDR = 3.0e-06)
NUPR1 (FDR = 1.4e-07)
GPNMB (FDR = 5.2e-09)
ALOX15B (FDR = 7.4e-07)
ZNF804A (FDR = 1.2e-05)
CXCR4 (FDR = 1.3e-04)
FPR3 (FDR = 1.2e-05)
TPRG1 (FDR = 1.1e-06)
GPR141 (FDR = 2.7e-04)
PLXNC1 (FDR = 1.6e-03)
counts per million
cluster
cl01 cl02 cl03 cl04 cl05 cl06 cl07 cl08 cl09 cl10 cl11 cl12 cl13
### cells
0
100
200
300
cluster status
other
test cluster
